# Deconvoluting the T cell response to SARS-CoV-2: specificity versus chance- and cognate cross-reactivity

**DOI:** 10.1101/2020.11.29.402677

**Authors:** Alexander A. Lehmann, Greg A. Kirchenbaum, Ting Zhang, Pedro A. Reche, Paul V. Lehmann

## Abstract

SARS-CoV-2 infection takes a mild or clinically inapparent course in the majority of humans who contract this virus. After such individuals have cleared the virus, only the detection of SARS-CoV-2-specific immunological memory can reveal the exposure, and hopefully the establishment of immune protection. With most viral infections, the presence of specific serum antibodies has provided a reliable biomarker for the exposure to the virus of interest. SARS-CoV-2 infection, however, does not reliably induce a durable antibody response, especially in sub-clinically infected individuals. Consequently, it is plausible for a recently infected individual to yield a false negative result within only a few months after exposure. Immunodiagnostic attention has therefore shifted to studies of specific T cell memory to SARS-CoV-2. Most reports published so far agree that a T cell response is engaged during SARS-CoV-2 infection, but they also state that in 20-81% of non-SARS-CoV-2-exposed individuals, T cells respond to SARS-CoV-2 antigens (mega peptide pools), allegedly due to T cell cross-reactivity with coronaviruses causing Common Cold (CCC), or other antigens. Here we show that by introducing irrelevant mega peptide pools as negative controls to account for chance cross-reactivity, and by establishing the antigen dose-response characteristic of the T cells, one can clearly discern between cognate T cell memory induced by SARS-CoV-2 infection vs. cross-reactive T cell responses in individuals who had not been infected with SARS-CoV-2.

## 1. Introduction

Traditionally, the assessment of immune memory has relied upon measurements of serum antibodies without queries of the T cell compartment. However, SARS-CoV-2 infection highlights the shortcoming of such a serodiagnostic approach. While the majority of SARS-CoV-2 infected individuals initially develop an antibody response to this virus, false negative results are a concern because not all infected individuals attain high levels of serum antibody reactivity acutely after infection ^1–3^, and those who do develop detectable antibody reactivity may decline to the limit of detection within a few months ^4^. In such cases, the detection of T cell memory might be the only evidence of such infection, and a surrogate of acquired immune protection from SARS-CoV-2 reinfection.

Fueled additionally by evidence that T cell-mediated immunity is required for immune protection against SARS-CoV-2 ^5–7^, attention has turned to T cell immunodiagnostics trying to establish whether the detection of T cell memory may be a more sensitive and reliable indicator of SARS-CoV-2 exposure than antibodies ^8–11^. In most studies published so far SARS-CoV-2-specific T cell memory cells were detected in the majority of infected individuals, but such were also found in 20-81% of control subjects who clearly could not have been infected by the SARS-CoV-2 virus ^9, 12–19^. If generalizable, such results would imply that T cell assays are unsuited to reliably identify who has, or has not, been infected by SARS-CoV-2 providing false positive results in up to 81% of the individuals tested. It should be noted right away, however, that the notion of cross-reactive SARS-CoV-2 antigen recognition by T cells being common in unexposed subjects might be related to the T cell assay and the test conditions used as it was not observed by others ^14, 20, 21^. Progress with settling the issue of T cell cross-reactivity in SARS-CoV-2 antigen recognition, and identifying suitable test systems, will decide whether T cell diagnostics can reliably detect specific immune memory to SARS-CoV-2 infection/exposure, and possibly identify the immune protected status of those subjects.

Next to possible cross-reactivity, T cell immune diagnostics of SARS-CoV-2 infection faces the challenge of having to reliably detect antigen-specific T cells in blood that occur in very low frequency. The numbers of SARS-CoV-2-specific T cells in blood post-infection is about one tenth of the numbers of T cells specific for viruses that induce strong T cell responses, such as influenza, Epstein Barr- (EBV) or human cytomegalovirus (HCMV) ^11, 22^, and reliably detecting even the latter is at the border of current technology. Further complicating matters, the frequencies of SARS-CoV-2-specific T cells are even lower in subjects who underwent a mild or asymptomatic SARS-CoV-2 infection compared to those who developed more severe COVID-19 ^12, 21, 23, 24^. Owing to these low T cell frequencies, and the antigen-induced signal being small in magnitude, any contribution of cross-reactive T cell stimulation will interfere with the reliable detection of genuine SARS-CoV-2-specific T cells. Setting up clear cut-off criteria for identifying antigen-specific T cell memory is therefore paramount.

Because T cell assays rely upon detecting SARS-CoV-2 antigen-specific T cells in blood via memory T cell re-activation *ex vivo,* the choice and formulation of the SARS-CoV-2 antigen itself used for the T cell recall will critically define the assay result. As the epitope utilization in the T cell response to SARS-CoV-2 is not known yet, by necessity, the aforementioned T cell diagnostic efforts tailored towards this virus have relied either on pools of hundreds of peptides that cover the entire proteome of the virus, or on pools of a multitude of predicted epitopes (mega peptide pools). Traditional T cell immune monitoring efforts, however, have called for the utilization of select, highly purified individual peptides whose specificity has been carefully established. Presently it is unproven whether pools of hundreds of unpurified peptides are even suited for reliable T cell diagnostics, and whether false positive or false negative results obtained using them are inherent to the recall antigen formulation. The chance for T cell cross-reactivity can be expected to increase with every peptide added into a pool, multiplying the chance for false positive results. Conversely, irrelevant peptides (those not recognized by T cells) also present in the pool can be expected to compete with the actually recognized T cell epitopes for binding to HLA-molecules, possibly causing false negative results ^25^. To our knowledge, it has not yet been systematically addressed whether and how chance cross-reactivity or peptide competition affects T cell immune monitoring results when mega peptide pools are used for testing. Instead of relying on third party mega peptide pools as the proper negative control to establish the background noise of the T cell assay, in all SARS-CoV-2 studies published so far, the mega peptide pool-induced T cell activation has been compared to PBMC cultured in medium alone, in the absence of any exogenously added peptide. In this report we introduce suitable negative control mega peptide pools, and using them, we address how to reliably detect even the very low frequency SARS-CoV-2 antigen-specific T cells in subjects who have undergone mild SARS-CoV-2 infection.

Cognate T cell cross-reactivity between related pathogens, such as SARS-CoV-2 and seasonal coronaviruses Common Cold (CCC), needs to be distinguished from the afore-mentioned chance cross-reactivities^19^. In cognate cross-reactivity, the TCR binds peptide sequences of two antigens that have extensive sequence homologies. In the majority of documented cases, such cross-reactive peptide sequences differ in only one or two amino acids with an additional requirement being that the exchange of amino acid(s) does not interfere with the peptides’ binding to, and folding in, the peptide binding groove of the restricting HLA molecule. Examples of experimentally verified cognate cross-reactivities include T cell recognition of serotypes of the Dengue virus ^26, 27^, influenza A virus strains ^28–31^, hepatitis C virus escape variants ^32^, and HIV epitope variants from different clades ^33, 34^.

It is still controversial how exclusively specific T cell antigen recognition is in general^35^. On one hand, there are reports suggesting that T cell recognition might be highly promiscuous with individual T cell clones being able to cross-reactively recognize 10^6^ different peptides ^36^. On the other hand, changing even a single amino acid in the presented peptide frequently abrogates T cell recognition, in particular if the change affects the binding of the peptide for the restricting MHC molecule, its conformation when bound to the MHC molecule, or when involving a TCR contact residue: while some studies have indicated an extremely low frequency of T cell cross-reactions between unrelated peptides ^37–39^, other studies (relying on tetramers) claim the opposite ^40, 41^. Accordingly, it needed to be addressed what impact TCR chance cross-reactivity has on *ex vivo* T cell monitoring when using mega peptide pools in general, and for SARS-CoV-2 antigen recognition in particular.

When T cell activation was seen in SARS-CoV-2-unexposed individuals using SARS-CoV-2 mega peptide pools for recall, the finding was interpreted as cognate cross-reactivity with related coronaviruses that cause harmless, common cold-like epidemies in the human population ^42^. There are four seasonal coronavirus strains, 229E, NL63, OC43, and HKU1, which cause pandemics in multiyear infection cycles in the human population world-wide ^43^. Although in any given year only 15-30% of humans displaying symptoms of common cold are indeed infected by one of these seasonal coronaviruses, 90% of the adult human population eventually becomes seropositive for at least three of these coronaviruses ^44–46^. From the perspective of T cell immune diagnostics of SARS-CoV-2, such cross-reactive T cell responses would generate false positive results. Another major scope of the present study was to establish to what extent cognate T cell cross-reactions of seasonal coronavirus antigens interferes with the detection of T cell memory induced by the SARS-CoV-2 virus itself.

The SARS-CoV-2 pandemic has made its rounds for nearly a year by now, yet its prevalence in the human population remains unknown as most of those infected go undiagnosed, having developed mild or no clinical symptoms at all ^47^. By now serum antibodies are no longer reliable in revealing, in retrospect, who has or has not been infected more than 3 months ago. If measurements of T cell memory would also fail to provide this information, our understanding of SARS-CoV-2’s prevalence will remain shrouded. Should vaccines under present development fail, without this information it will remain guesswork to decide whether and when sufficient herd immunity has developed in a population, or if robust immunity develops at all following natural infection ^48^. Without knowing who has or has not been infected by SARS-CoV-2, one cannot distinguish whether a candidate vaccine can prime a protective immune response in naïve individuals, or whether it merely boosts immunity that has been pre-established by the natural infection. Without this information, all those individuals – possibly the majority of the population - who already went through an uncomplicated SARS-CoV-2 infection and might be protected from re-infection, or are prone to develop a mild disease if reinfected again, need to continue to live in fear of contracting a potentially lethal disease.

In this report we sought solutions to deconvolute T cell reactivity to SARS-CoV-2 mega peptide pools so as to clearly distinguish between individuals who have or have not been infected with this virus.

## 2. Materials and Methods

### 2.1 Peripheral Blood Mononuclear Cells

Pre-COVID Era Donors. PBMC from healthy human donors were obtained from CTL’s ePBMC library (CTL, Shaker Heights, OH, USA) collected prior to Dec 31, 2019. The PBMC had been collected by HemaCare Blood Donor Center (Van Nuys, CA) under HemaCare’s IRB and sold to CTL identifying donors by code only while concealing the subjects’ identities. All PBMC were from healthy adults who had not taken medication within a month of the blood draw that might influence their T cell response. In addition, tests were done on each donor at HemaCare’s CLIA certified laboratory to identify common infections, including Human Immunodeficiency Virus (HIV). Subjects positive for HIV were disqualified for the ePBMC library. The donors’ age, sex, and ethnicity are shown in S. Table 1.

SARS-CoV-2 Infected Donors. PBMC of subjects were collected under Advarra IRB Approved # Pro00043178, CTL study number: GL20-16 entitled COVID 19 Immune Response Evaluation. All subjects tested positive for SARS-CoV-2 RNA in PCR performed on nasal swabs, and these tests were performed in accredited medical laboratories. All such donors underwent mild COVID infection from which the subjects fully recovered within one or two weeks. These subjects were bled between 2 weeks and 3 months post recovery (median of 24 days). The donors’ age, sex, and ethnicity are shown in S. Table 1. PBMC were isolated and cryopreserved at CTL.

The cryopreserved cells were thawed following an optimized protocol ^49^ resulting in viability exceeding 90% for all samples. The PBMC were resuspended in CTL-Test^TM^ Medium (from CTL). CTL-Test^TM^ Medium is serum-free and has been developed for low background and high signal performance in ELISPOT assays. The number of PBMC plated in the ELISPOT experiments was 2 × 10^5^ viable PBMC per well.

### 2.2. ELISA Assays

MaxiSorp 96-well microplates (Thermo Fisher) were coated with recombinant SARS-CoV-2 Nucleocapsid (RayBiotech, Peachtree Corners, GA), truncated Spike protein (S1 domain) (The Native Antigen Company, Oxford, UK) or receptor binding domain (RBD) (Center for Vaccines and Immunology (CVI), UGA, Athens, GA) at 2ug/mL in PBS overnight at 4°C. Plates were then blocked with ELISA blocking buffer containing 2% w/v bovine serum albumin in PBS with 0.1% v/v Tween20 (PBS-T) (Sigma-Aldrich) for 1 h at room temperature. Donor plasma were serially diluted in assay plates and incubated overnight at 4°C. Plates were then washed with PBS prior to addition of horseradish peroxidase-conjugated anti-human IgG detection reagents (from CTL) and incubation for 2 h at room temperature. Plates were then washed with PBS prior to development with TMB chromogen solution (Thermo Fisher). 1M HCl was used to stop conversion of TMB and optical density was measured at 450nm (OD_450_) and 540nm (OD_540_) using a Spectra Max 190 plate reader (Molecular Devices, San Jose, CA USA). Optical imperfections in assay plates were corrected through subtraction of OD540 values. Antigen-specific IgG concentrations are reported as μg/mL IgG equivalents and were interpolated from a standard curve generated using an IgG reference protein (Athens Research and Technology, Athens, GA) coated directly into designated wells of assay plates.

### 2.3. Antigens and Peptides

All mega peptide pools used in this study are products of, and were purchased from JPT (Berlin, Germany). The peptide pools representing the individual antigens are shown in S. Table 2. All these mega peptide pools consisted of 15-mer peptides that covered the entire amino acid (aa) sequence of the respective proteins in steps (gaps of) 11 aa. All mega peptide pools were tested at a final concentration of 1.5 μg/mL of each peptide within the pool at the highest concentration, followed by three 1+2 (vol + vol) serial dilutions, as specified in the Tables. CPI (from CTL) was used as a positive control for the activation of antigen-specific CD4+ T cells ^50^. CPI is a combination of protein antigens derived from CMV, influenza and parainfluenza viruses, and was used at a final concentration of 6.25 μg/mL in ELISPOT assays. CERI (from CTL) was used as a positive control for CD8+ T cells ^51^. CERI is a mega peptide pool consisting of 124 peptides of <u>C</u>ytomegalo- (CMV), <u>E</u>pstein-Barr-(EBV), Human <u>r</u>espiratory syncytial-(HRSV) and <u>I</u>nfluenza viruses. The individual peptides, 9 amino acids long, were selected based upon peptide binding predictions for a broad range of HLA class I alleles expressed in all human races and diverse ethnic subpopulations. Both CPI and CERI elicit T cell recall responses in all healthy donors tested so far^51^.

All mega peptide pools were delivered as lyophilized powder. The individual peptide pools were initially dissolved following the manufacturer’s directions in 40 μl DMSO, followed by addition of 210 μl of PBS generating a “primary peptide stock solution” at 100 ug/mL (0.1mg/mL) with 16% v/v DMSO. From each of these wells, a “secondary peptide stock solution” was prepared in a 96-Well Deep Well Plate, with peptides starting at 3 ug/mL which were then threefold serially diluted. Using a 96-well multichannel pipettor 100ul was transferred “en block” into a pre-coated ImmunoSpot^®^ assay plates. Finally, 100 μL of PBMC (containing 2 × 10^5^ cells) in CTL-Test media was added “en block” to achieve the desired final peptide concentrations of 1.5, 0.5, 0.17 and 0.06 ug/mL in the ELISPOT assay.

### 2.4. Human IFN-γ ELISPOT Assays

Single-color enzymatic ImmunoSpot^®^ kits from CTL were used for the detection of in vivo-primed IFN-γ-producing Th1-type memory T cells. Test procedures followed the manufacturer’s recommendations. In brief, peptides were plated at the specified concentrations into capture antibody-precoated ELISPOT assay plates in a volume of 100 μL per well, dissolved in CTL-Test™ Medium. The plates with the antigen were stored at 37°C in a CO_2_ incubator for less than an hour until the freshly thawed PBMC were ready for plating. The PBMC were added at 200,000 viable cells /well in 100 μL CTL-Test™ Medium and cultured with the peptides for 24h at 37 °C and 9% CO_2_ in an incubator. After removal of the cells, addition of detection antibody, and enzymatic visualization of plate-bound cytokine, the plates were air-dried prior to scanning and counting of spot forming units (SFU). ELISPOT plates were analyzed using an ImmunoSpot^®^ S6 Ultimate Reader, by CTL. SFU numbers were automatically calculated by the ImmunoSpot^®^ Software for each stimulation condition using the Autogate^TM^ function of the ImmunoSpot^®^ Software ^52^.

### 2.5. Statistical Analysis

As ELISPOT counts follow Gaussian (normal) distribution among replicate wells, the use of parametric statistics was justified to identify positive and negative responses, respectively. Positive responses were defined as SFU counts exceeding 3 SD of the mean SFU counts of the specified negative control, identifying such at 99.7% confidence.

## 3. Results and Discussion

### 3.1. SARS-CoV-2-Antibody Reactivity in Subjects Recovered from Mild COVID-19

The primary goal of this study was to characterize T cell memory to SARS-CoV-2 antigens in subjects who recovered from PCR-verified mild COVID-19 infection. We gained access to nine such subjects’ blood, referring to them as “SARS-CoV-2 PCR-Positive Subjects” in this communication. First, we tested whether these individuals exhibited evidence of IgG reactivity against SARS-CoV-2 antigens. As shown in Figure 1A, and consistent with prior findings ^53, 54^, IgG reactivity against the SARS-CoV-2 Nucleocapsid protein was significantly (p<0.01) increased in these subjects compared to our control cohort, individuals who were bled in the pre-COVID-19 era. We refer to these latter donors as “Pre-SARS-CoV-2 Era Subjects”. Similarly, IgG reactivity against the truncated Spike (S1) protein was also significantly (p<0.001) elevated in the COVID-19-recovered donors (Figure 1B) ^55^. Despite PCR confirmation of recent SARS-CoV-2 infection, 1/3 of the COVID-19 recovered donors exhibited indistinguishable IgG binding to the Nucleocapsid and S1 probes compared to the Pre-COVID-19 collected samples. Collectively, these binding assays demonstrate that measurement of serum IgG reactivity against the Nucleocapsid and/or S1 proteins alone is insufficient to identify all donors who were recently infected with the SARS-CoV-2 virus.

**Figure 1.**
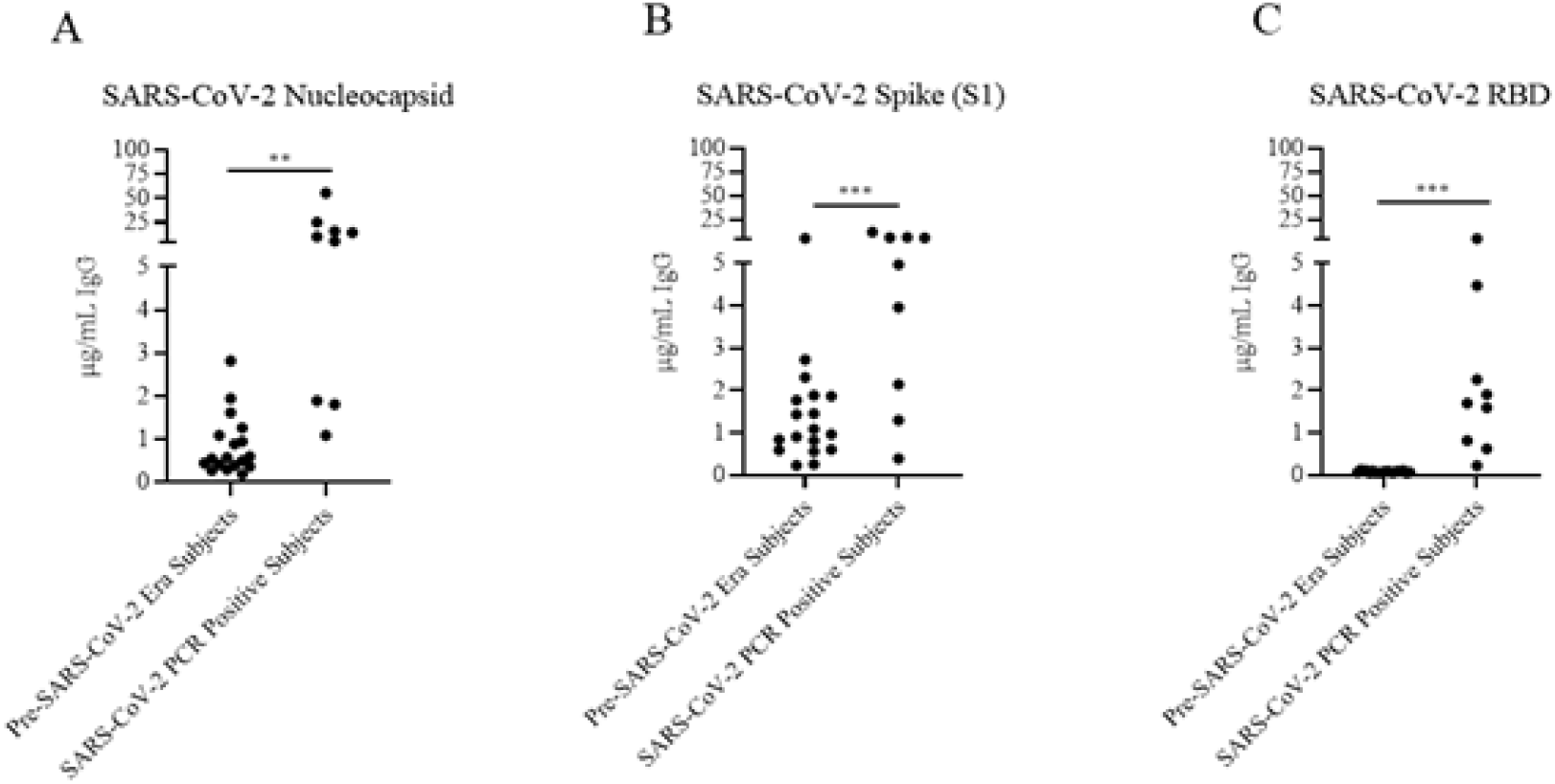
IgG antibody reactivity against SARS-CoV-2 antigens. Plasma from 18 Pre-SARS-CoV-2 Era Subjects and 9 individuals who underwent mild PCR-confirmed SARS-CoV-2 infection (SARS-CoV-2 PCR-Positive Subjects) were assessed for IgG antibody against A) SARS-CoV-2 Nucleocapsid protein, B) the truncated SARS-CoV-2 Spike fragment, S1, or C) the Spike RBD fragment. ELISA binding signal was then interpolated into ug/mL IgG equivalents using a reference standard. Each serum sample is represented by a dot. Statistical significance between the two cohorts was determined using an unpaired Student’s t-test. Significant differences between cohorts are marked with ** denoting p < 0.01, and *** p < 0.001, respectively.

In contrast to the Nucleocapsid and S1 probes, IgG reactivity against the receptor binding domain (RBD) of the SARS-CoV-2’s Spike protein was not detected in the Pre-SARS-CoV-2 Era Subjects (Figure 1C), but was significantly (p<0.001) increased in the SARS-CoV-2-PCR Positive Cohort. Thus, the presence of IgG reactivity against the RBD probe appears to be a reliable indicator of recent virus infection) ^55^, and possibly also of neutralizing potency ^55, 56^.

Two of 9 donors in our COVID-19-recovered cohort exhibited low IgG reactivity against each of the SARS-CoV-2 probes. (These were subjects dC4 and dC7, of whom dC4 did reveal T cell memory, but dC7 did not, see below, Table 2). In the absence of the PCR-confirmed diagnosis, their respective exposures to SARS-CoV-2 would be difficult to verify if solely based on serologic assessment. Our observations are consistent with the notion that the severity of clinical disease and the magnitude of the elicited humoral response to SARS-CoV-2 are positively correlated ^4, 55^. Weak antibody responses in sub-clinically infected subjects, along with recent evidence for declining antibody titers within the first 3 months of convalescence ^4^, pose a substantial challenge to serology-based assessment of SARS-CoV-2 immunity. Hence, we focused on the detection of T cell memory to this virus.

### 3.2. Experimental Design for Assessment of T Cell Memory to SARS-CoV-2

#### 3.2.1. The Rationale for Selecting IFN-γ ELISPOT for Detecting SARS-CoV-2-Specific Memory T cells

T cell immune monitoring aims at detecting *in vivo* expanded and differentiated antigen-specific T cell populations directly *ex vivo*, either in freshly isolated PBMC, or in PBMC that have been cryopreserved following protocols that maintain full T cell functionality upon thawing the cells ^57^. The number (frequency) and functions (e.g. cytokine signature) of antigen/peptide-specific T cells need to be measured as present in the body without inducing additional clonal expansions or T cell differentiation *in vitro* during the short-term *ex vivo* antigen stimulation that is required to detect the antigen-reactive T cells. In ELISPOT/ImmunoSpot^®^ assays, antigen- (peptide)-specific T cells present in the PBMC become activated, and they start producing cytokine. This cytokine is captured on a membrane around each secreting T cell, resulting in a cytokine spot (a spot forming unit, SFU). Counting of SFUs permits to establish, at single-cell resolution, the number of antigen-triggered cytokine-producing T cells ^58^, and thus the frequency of such cells in PBMC. In this study we focused on IFN-γ measurements because, in subjects who successfully overcome SARS-CoV-2 infection, Th1 cells have been reported to prevail by far ^5, 9, 18, 59, 60^. Th1 cells have been implicated as a protective class of response while Th2 and Th17 have been linked to immune pathology ^59, 61^. Furthermore, the standard 24h IFN-γ ELISPOT assay detects *in vivo* primed Th1 effector memory cells only; naïve T cells or central memory cells are not detected in this assay as the latter require several days of differentiation following antigen encounter before they begin secreting IFN-γ ^62, 63^.

We also selected the *ex vivo* ELISPOT platform because it requires as few as 200,000 peripheral blood mononuclear cells (PBMC) per antigen stimulation condition, and this assay lends itself to high throughput analysis. Utilizing only 32 million PBMC per subject (obtainable from 32 mL of blood), we established 155 T cell reactivity datapoints per subject, testing 37 mega peptide pools (see S Table 2), each at four concentrations, plus 4 media and 3 positive control wells, all in a single high-throughput experiment. In the mega peptide pools, the individual peptides were present at 1.5 μg/mL at the highest concentration tested, and in 0.5 μg/mL, 0.17 μg/mL, and 0.06 μg/mL in the subsequent 1 + 2 (vol+vol, 3-fold) serial dilutions. Instead of using replicates, these serial antigen dilutions served not only to confirm positive results, but additionally permitted us to establish the affinity of the responding T cells.

#### 3.2.2. The Rationale for Using Mega Peptide Pools for Detecting SARS-CoV-2-Specific Memory T cells

Due to the highly individualized nature of T cell epitope recognition in general, for which evidence is also starting to accumulate for SARS-CoV-2 ^11–13, 16, 21, 60^, and due to the size of the virus (whose genome is approximately 29.8 kb ^64^ there are only two viable options for selecting peptides for a comprehensive assessment of T cell immunity to SARS-CoV-2. One option is to perform in silico epitope predictions, and such has already been reported for SARS-CoV-2 ^65^, but their accuracy has recently been convincingly called into question ^66, 67^. Moreover, for a comprehensive assessment, the epitope predictions would need to be customized for each test subject, accounting for all HLA class I and class II molecules expressed in each individual. Because it is impractical to individualize predicted peptide epitopes for each subject, we have elected to take the agnostic route, in which the entire sequence of each protein antigen is covered by a series of overlapping peptides. However, this means that, dependent upon the length of the protein, hundreds of peptides need to be combined into mega pools (see S. Table 2). While, in theory, the mega peptide pool approach permits systematic coverage of all possible T cell epitopes within a virus, it introduces an as yet undefined dimension: chance cross-reactivity between unrelated peptides.

#### 3.2.3. The Rationale for Selecting Negative Control Mega Peptide Pools

To account for chance T cell cross-reactivity, we tested mega peptide pools covering foreign antigens to which it is unlikely the test subjects have been exposed (one Ebola virus peptide pool, five HIV antigen pools) and a self-antigen, Actin, that due to its abundance in the body is likely to have established self-tolerance. These mega peptide pools are defined in S. Table 2, including the number of peptides contained in each.

#### 3.2.4. Avoiding Inter-Assay Variations

To reduce assay variables, all peptide pools used in this study were from the same vendor, were synthetized, stored, dissolved and tested the same way, and had the exact same formulation consisting of 15 amino acid long peptides that systematically walk the entire sequence of the respective proteins in steps of 11 amino acids. Taking advantage of the high-throughput suitability of ELISPOT, all the peptide pools (S. Table 2) and their dilutions were tested on each PBMC donor in a single experiment which rendered the peptides the only assay dependent variable. This approach therefore permitted us to firmly establish within each PBMC sample the number of Th1 T cells responding to the different mega peptide pools, and thus to compare the frequencies of the respective mega peptide pool-reactive T cells within each PBMC donor, and amongst donors in the cohorts.

We compared T cell reactivity to all the above mega peptide pools in 18 healthy Pre-COVID-19 Era Subjects and in the 9 individuals who recovered from mild SARS-CoV-2 infection as verified by PCR. In the following we describe and interpret the results.

### 3.3. Classic Single Antigen and Single Antigen Dose-Based Data Analysis does not Permit to Distinguish Between SARS-CoV-2-Infected Subjects and Controls

Figure 2 shows the test results comparing frequencies of SARS-CoV-2 antigen-reactive IFN-γ-producing T cells in the SARS-CoV-2 PCR-verified (also referred to as COVID-recovered) and Pre-SARS-CoV-2 (hereafter referred to as Pre-COVID) cohorts when tested at a single antigen concentration, 1.5 μg/mL of peptide within each peptide pool. Essentially identical results have been reported by others ^9, 12–18^ showing that, as a cohort, the frequency of SARS-CoV-2 antigen-specific T cells is significantly elevated in COVID-recovered individuals versus the cohort that has not been infected with the SARS-CoV-2 virus. In all these aforementioned studies, and for T cell immune monitoring in general, it has been of concern however, that such results are inconclusive for the individuals, as a large fraction of COVID-recovered subjects show similar or even lower frequencies of SARS-CoV-2 antigen-specific T cells than the Pre-COVID control subjects. Simple T cell frequency measurements using single SARS-CoV-2 antigens at a single antigen concentration therefore do not permit to reliably distinguish whether an individual has or has not been infected by SARS-CoV-2. This finding, along with the low frequency of SARS-CoV-2 antigen-reactive T cells, may come as a surprise because massive clonal expansions are typically seen initially after infections and vaccinations ^63^. There is increasing evidence that the SARS-CoV-2 virus actively disrupts the engagement of an immune response ^68–74^ explaining its weak immunogenicity. The low frequencies of SARS-CoV-2 antigen-reactive T cells in COVID-recovered individuals in turn makes it challenging to unambiguously detect them.

**Figure 2.**
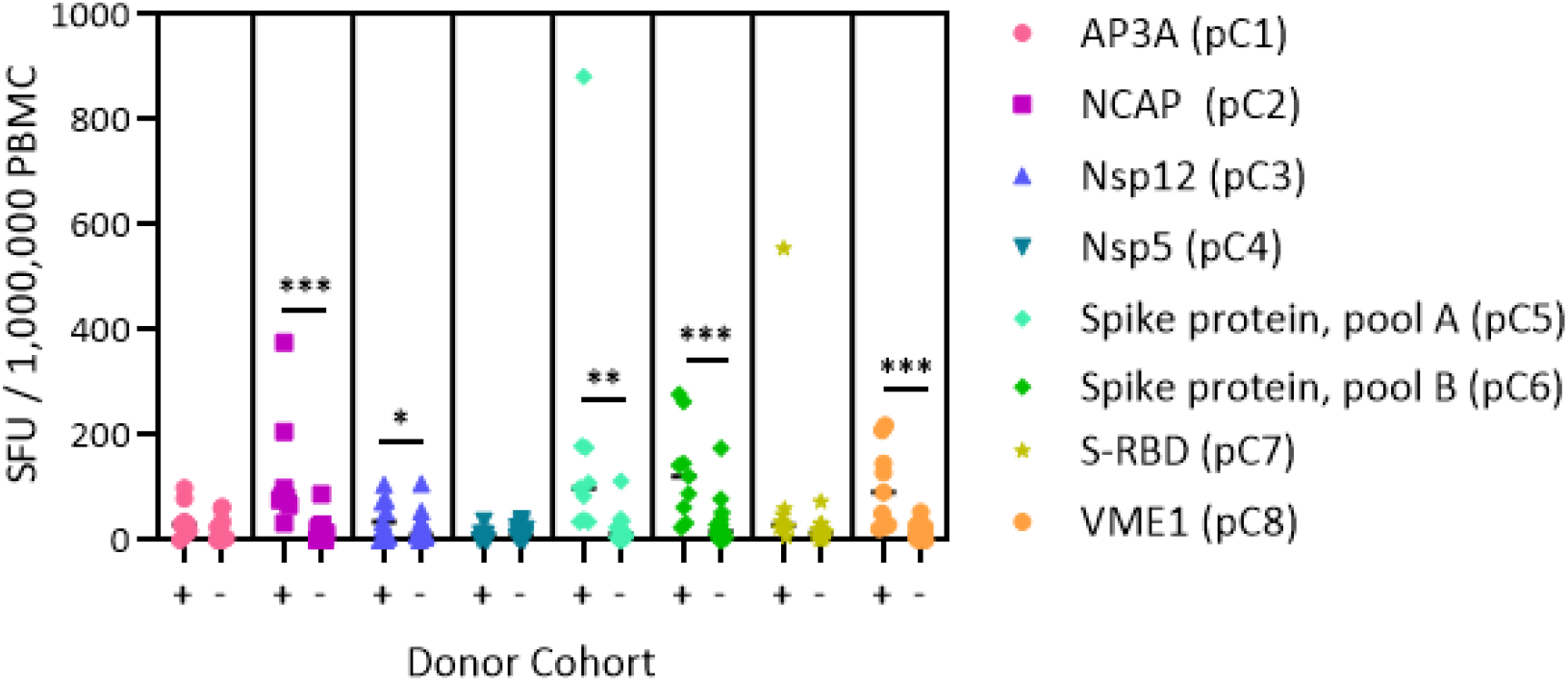
Classic representation of SARS-CoV-2 antigen-specific T cell frequencies in SARS-CoV-2 PCR-verified Subjects (+) versus Pre-Covid Era Subjects (−) cohorts. PBMC of each individual within the cohort is represented by a dot. The PBMC were challenged with the SARS-CoV-2 peptide pools specified on the right, each pool covering different antigens of the virus (see Suppl. Table 2). The individual peptides in each pool were tested at 1.5 μg/mL. An ELISPOT assay was performed measuring the numbers of antigen-induced IFN-γ-secreting cells, spot forming units (SFU), in 200,000 PBMC; following convention, the numbers have been normalized to per million PBMC, as shown on the Y axis. Significant differences between SARS-CoV-2-infected vs. non-exposed cohorts are marked with * denoting p ≤ 0.05, ** p ≤ 0.01, and *** p ≤ 0.001, respectively.

### 3.4. Accounting for Chance Cross-Reactivity when Testing Suitable Control Mega Peptide Pools to Establish the Background Noise in ELISPOT Assays

The challenge with selecting mega peptide pools that are suited for negative controls (PP. Neg Contr.) is that one needs to identify antigens to which the test population has not been exposed. As one such antigen we selected the Ebola virus nucleoprotein (pN6, standing for peptide pool negative control 6; abbreviations used for peptide pools and the specifics of them are listed in S. Table 2). We also included five HIV antigens (nN2, pN3, Pn4 and pN7) as the participating subjects needed to be HIV-seronegative to qualify for this study. Finally, we tested as a negative control a peptide pool that covers the sequence of the self-antigen, Actin (pN5). As we did for SARS-CoV-2 peptide pools, all these PP. Neg Contr. candidates were tested at four concentrations, 1.5 μg/mL, 0.5 μg/mL, 0.17 μg/mL, and 0.06 μg/mL of each peptide within the pool. The number of PP. Neg Contr. candidate-induced IFN-γ-producing cells was compared to the medium control, the latter measured in quadruplicate wells. CPI, CERI, and CEFX antigens were measured in singlet, and served as positive controls, respectively ^51^. For each PBMC sample, the mean and standard deviation (SD) of four replicate medium control wells was established, and compared to the SFU counts induced by the candidate negative control peptide pools. SFU counts greater than 3 SD of the mean medium control counts are highlighted in Table 1.

**Table 1.**
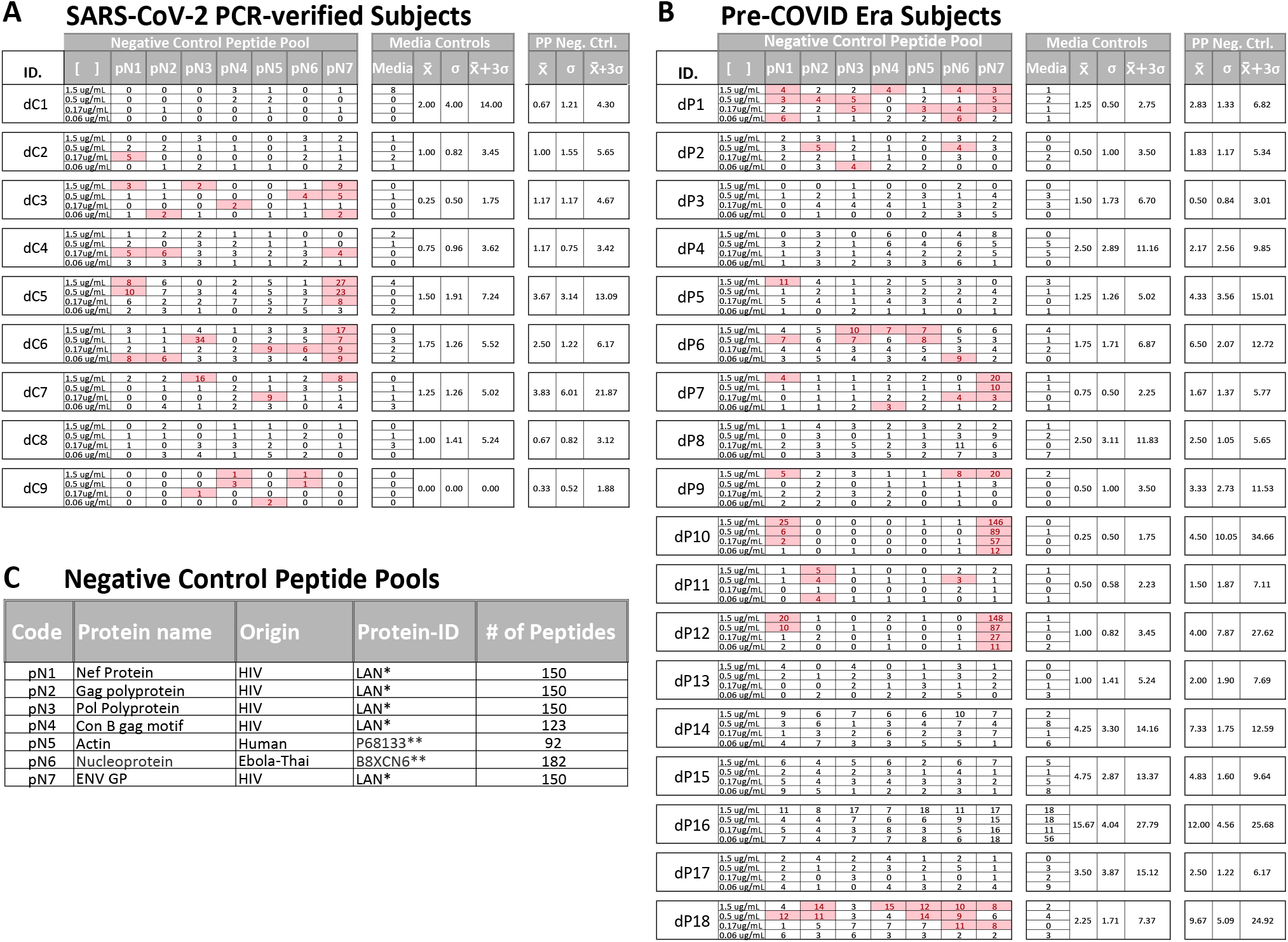
IFN-γ SFU counts triggered in both cohorts by control peptide pools. PBMC of 9 subjects with SARS-CoV-2-PCR-verified infection (A) and PBMC from 18 subjects from Pre-Covid Era (B) where tested in an ELISPOT assay with the peptides specified in (C). These peptide pools were tested in 4 serial dilutions on each PBMC sample. Highlighted are SFU counts that exceeded the mean of the four medium control SFU counts by more than 3 SD for the respective subject, as listed under “Media Control”. The mean, SD and 3 SD for the Peptide Negative Control (PP Neg. Cont.) is also shown for each subject as calculated from the SFU counts triggered by the six negative control peptide pools, pP1-pP6, at 1.5 ug/mL, with the exclusion of pN7 (HIV Core protein), for reasons specified in the Text.

The peptide pool covering HIV Envelope (Env) protein (pN7) induced vigorous SFU formation (> 100 SFU/200,000 PBMC) in 3 subjects’ PBMC, and relatively strong SFU formation (17-42 SFU/200,000 PBMC) in two additional subjects, recalling positive responses in 3-4 consecutive peptide dilutions. At the highest concentration, this peptide pool also elicited elevated SFU numbers in 4 additional subjects. All these subjects who responded to HIV Env protein were HIV-seronegative as established by the blood bank (Hemacare, Van Nuis, CA) that collected them. The HIV Env protein belongs to the p24 superfamily, which is shared sequence conservation with related proteins expressed by many retroviruses ^75^. Thus, cognate cross-reactivity with T cells primed by such retroviruses strikes us as the likely explanation for the pN7-triggered recall responses seen in HIV-seronegative subjects. Be that as it may, the HIV Env peptide pool is clearly unsuited as a negative control peptide pool.

The six remaining candidate negative control peptide pools occasionally triggered elevated SFU counts, but these occurred at relatively low frequencies, and predominantly at only the highest peptide concentration (Table 1). The data therefore provide evidence for low-level chance cross-reactivity when mega peptide pools are tested in ELISPOT assays. In theory, at higher peptide concentrations this chance cross-reactivity might increase, however, as T cells with low affinity for the peptides also reach their activation threshold.

Therefore, when analyzing the following SARS-CoV-2 peptide pool-triggered T cell responses, we will use – and compare – two negative controls. One is the conventional “Medium control”, established as the mean and SD of 4 replicate wells in which PBMC were cultured with medium alone. The second is the “peptide pool negative control” (PP Neg. Control.), calculated as the mean and SD of each donor’s SFU count induced by the six negative control peptide pools at 1.5 μg/mL. Again, these six negative control pools encompassed Ebola, Actin, and the 4 HIV antigen pools (excluding HIV Env). The mean and SD for the PP Neg. Control, and Medium control, and the raw data from which these were derived are specified for each subject in Table 1.

### 3.5. Chance Cross-Reactivity Accounts for Most of SARS-CoV-2 Peptide Reactivity in Pre-COVID Era Subjects

We compared the SFU counts induced by SARS-CoV-2 mega peptide pools in the subjects who recovered from mild COVID-19, and those who were bled prior to the COVID era. The SARS-CoV-2 peptide-induced SFU counts were analyzed vs. either the Medium control, or the PP. Neg. Control, in each case highlighting positive SFU counts as defined by exceeding 3 SD of the respective mean control count, a threshold that identifies positive responses with > 99.6% confidence. As can be seen in S. Table 3, in Pre-COVID subjects the number of SARS-CoV-2 peptide-induced positive SFU counts was significantly lower when control peptide pools were used as to establish the background noise level. Thus, chance cross-reactivity, rather than cognate cross-reactivity with seasonal coronaviruses, accounted for most of the positive responses detected in Pre-COVID control subjects. The few apparently positive cross-reactive responses left after filtering for chance cross-reactivity in this cohort could be discerned from cognate T cell responses to SARS-CoV-2 peptides in SARS-CoV-2 PCR-verified individuals when taking the affinity of the T cell response into account, as will be shown below.

### 3.6. Affinity for SARS-CoV-2 Peptides Distinguishes Cognate from Cross-Reactive T cell Recognition

Testing peptide pools in four serial dilutions not only permits generation of confirmatory results without using replicate wells, but also permits one to gain insights into the affinity of the T cells recognizing the respective peptides. In the context of this study, we will distinguish between Level 4 affinity (high affinity, with all four peptide concentrations recalling T cells, color coded in red), Level 3 affinity (intermediate affinity, eliciting a recall response across 3 consecutive peptide dilutions, highlighted in orange), Level 2 affinity (low affinity, only the two highest peptide concentrations elicit a T cell response, color coded in yellow), and Level 1 affinity (borderline low, eliciting a significant T cell response at the highest peptide concentration only, color coded in beige).

As seen in Table 2, most COVID-recovered subjects displayed Level 4 (red) affinity T cell responses to several SARS-CoV-2 antigens, while this level was absent in the Pre-COVID Era controls. In the latter, only occasional Level 3 (orange) and Level 2 affinities (yellow) were seen. Thus, high affinity responses to several SARS-Cov-2 antigens (unlike responses detected against individual antigens at a single antigen concentration, see Fig. 1) appear to be suited to distinguish cognate SARS-CoV-2 specific T cells in COVID-recovered subjects from cross-reactive T cells in subjects infected by other coronaviruses in the Pre-COVID Era. The frequency of SARS-CoV-2 mega peptide pool-specific T cells was low, but clearly elevated > 3 SD over the negative control mega peptide pool control level. As ELISPOT SFU counts follow normal distribution ^76^, the mean of background plus ≥ 3 SD positivity cut-off definition sets the chances for a single datapoint being a false positive at ≤0.4%; for four responses in a row being false positive, the chances are negligible at a probability of < 0.0256%.

**Table 2.**
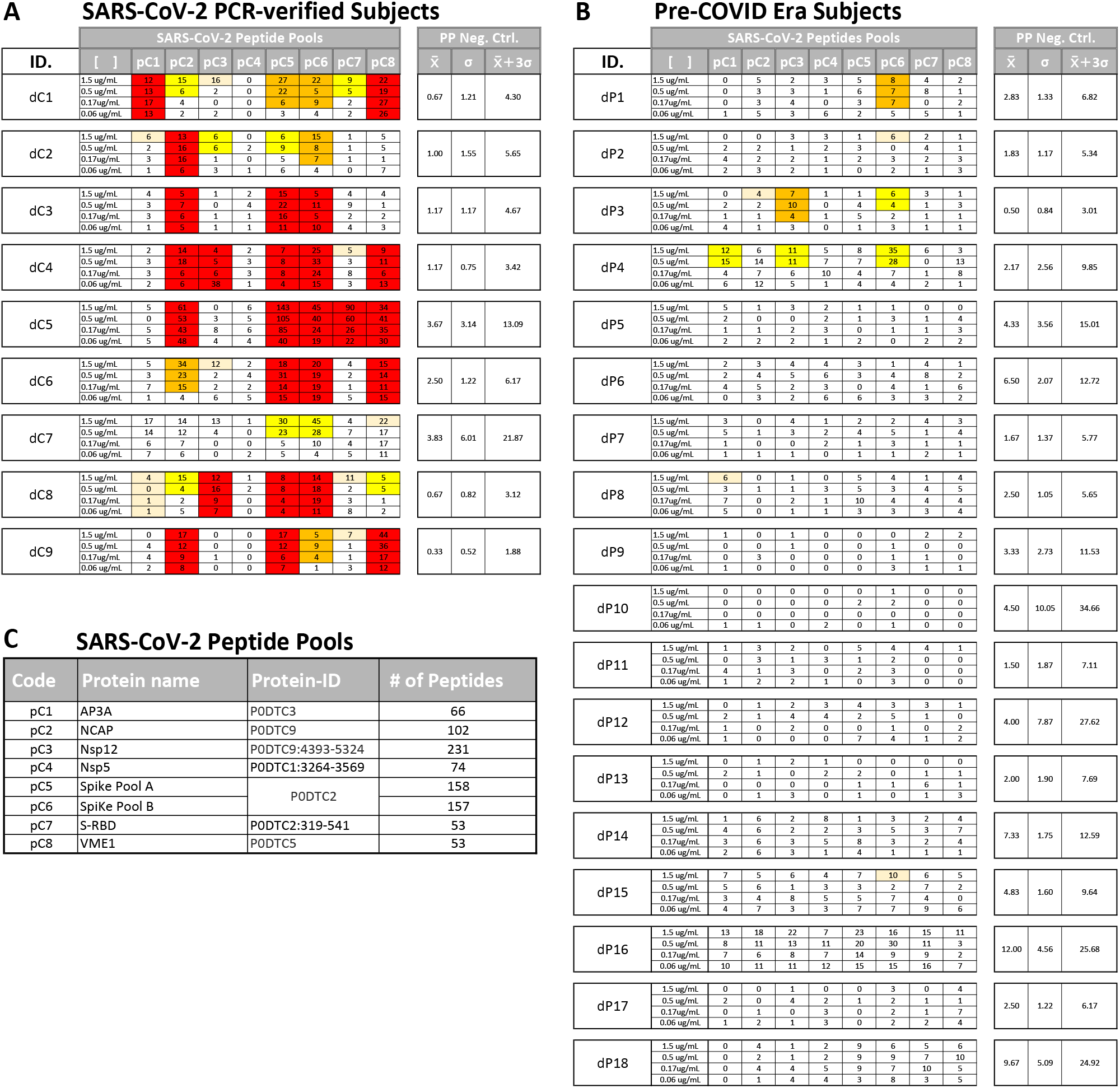
Affinity analysis of SFU counts triggered by SARS-CoV-2 peptides in PBMC of donors with SARS-CoV-2-PCR-verified infection (A) and in pre-Covid Era subjects (B). Affinity levels are color-coded. Red: high affinity, defined as four consecutive peptide dilutions eliciting a positive recall response with SFU counts exceeding 3 SD of the negative peptide pool-based background (PP Neg. Contr.), as specified for each PBMC sample on the right. Orange: intermediate affinity, defined as three consecutive peptide dilutions eliciting a positive recall response. Yellow: low affinity, with only the two highest peptide concentrations eliciting positive SFU counts.

These types of affinity measurements, which rely on serial dilution of peptides and are simple to perform, also require high-throughput suitable test platforms that are frugal with regards PBMC utilization, such as ELISPOT. To our knowledge, such T cell affinity measurements have so far not been applied systematically to characterize virus-specific T cell responses, thus permitting to compare the above affinity distributions observed for SARS-CoV-2 with other viruses. To compare the SARS-CoV-2 antigen-induced T cell responses with T cell reactivity to a better characterized virus, we also tested mega peptide pools that covered 14 antigens of Epstein Barr Virus (EBV), which commonly infects most humans by the time they have reached adulthood. The raw data are shown in S. Table 4. As summarized in Table 3, the percentage of EBV mega peptide pools recognized at affinity Levels 1-4 was comparable in both cohorts to the percentage of SARS-CoV-2 peptide pools eliciting Level 1-4 T cell recall responses in the COVID-recovered subjects. SARS-CoV-2 infection, therefore, seems to induce a T cell response that, at least as far as the affinity of nominal antigen-recognition goes, is comparable to the T cell response to EBV.

**Table 3.** Affinity distributions of T cells recognizing SARS-CoV-2 peptides in SARS-CoV-2-recovered and Pre-COVID era subjects (A) vs. the affinity distribution of T cells recognizing EBV peptides in both cohorts. Peptide pools eliciting positive T cell recall responses in the specified affinity level categories within the cohort are shown in absolute numbers (No. Positive) or as the percentage of all positive responses within the cohort (% Positive). The raw data are shown in Table 2 for the SARS-Cov-2 peptides, and in S. Table 5 for the EBV peptides.

**A**
| Aff. Level | Definition | Color Code | SARS-CoV-2 Peptide Pools Positive (%) |  |
| --- | --- | --- | --- | --- |
|  |  |  | COVID-recovered Subjects | Pre-COVID Era Subjects |
| 4 | 4 serial + peptides |  | 31% | 0% |
| 3 | 3 serial + peptides |  | 6% | 3% |
| 2 | 2 serial + peptides |  | 10% | 5% |
| 1 | First + only |  | 10% | 5% |

**B**
| Aff. Level | Definition | Color Code | EBV Peptide Pools Positive (%) |  |
| --- | --- | --- | --- | --- |
|  |  |  | COVID-recovered Subjects | Pre-COVID Era Subjects |
| 4 | 4 serial + peptides |  | 28% | 36% |
| 3 | 3 serial + peptides |  | 6% | 12% |
| 2 | 2 serial + peptides |  | 6% | 11% |
| 1 | First + only |  | 9% | 23% |

### 3.7. Fine Specificity of SARS-CoV-2 Antigen Recognition in COVID-19-Recovered Subjects

The SARS-CoV-2 peptide pools we used in our study encompassed eight major viral proteins, and systematically covered the respective antigens. Comparing within each donor the SFU counts triggered by these peptide pools permits therefore to assess, first, the total T cell mass mobilized against the virus, and second which antigens are preferentially targeted by T cells, i.e., the T cell immune dominance hierarchy within SARS-CoV-2 antigens. As shown in Table 4, Spike protein (pC5&6) was dominant, or co-dominant, in all COVID-recovered subjects, with 24-51% of all SARS-CoV-2-specific T cells targeting this protein in the individual subjects (it was 40±9% for the cohort). The recognition of Nucleocapsid(NCAP) (pC2, 18±6%) and VMA-1 (pC8, 16±12%) was next most abundant for the COVID-recovered cohort, while S-RBD (pC7, 9±6%), Nsp12 (pC3, 8±6%) and AP3A (pC1, 6±4%) peptide pools constituted third tier targets for T cells. There was therefore a clear T cell response hierarchy at the level of the cohort, but it did not always hold up for each individual within the cohort. For subject dC9, for example, only 24% of the SARS-CoV-2-specific T cells targeted Spike protein vs. 49% being specific for VMA-1, 19% for NCAP, and 8% targeting S-RBD. An immune monitoring effort that focused only on the “immune dominant” Spike protein would have detected only 24% of the relevant T cells in this subject. In Subject dC2, 40% of the SARS-CoV-2-specific T cells targeted Spike protein, but these T cells were of lower affinity than the 25% that recognized NCAP. T cell immune monitoring efforts for SARS-CoV-2 therefore ideally should include all antigens of the virus, tested in serial dilutions. Supplemental Table 4 shows for EBV how inaccurate the assessment of T cell immunity to this virus would be if it was restricted to a single antigen, and at a single peptide dose. The same holds for HCMV ^77^.

**Table 4.**
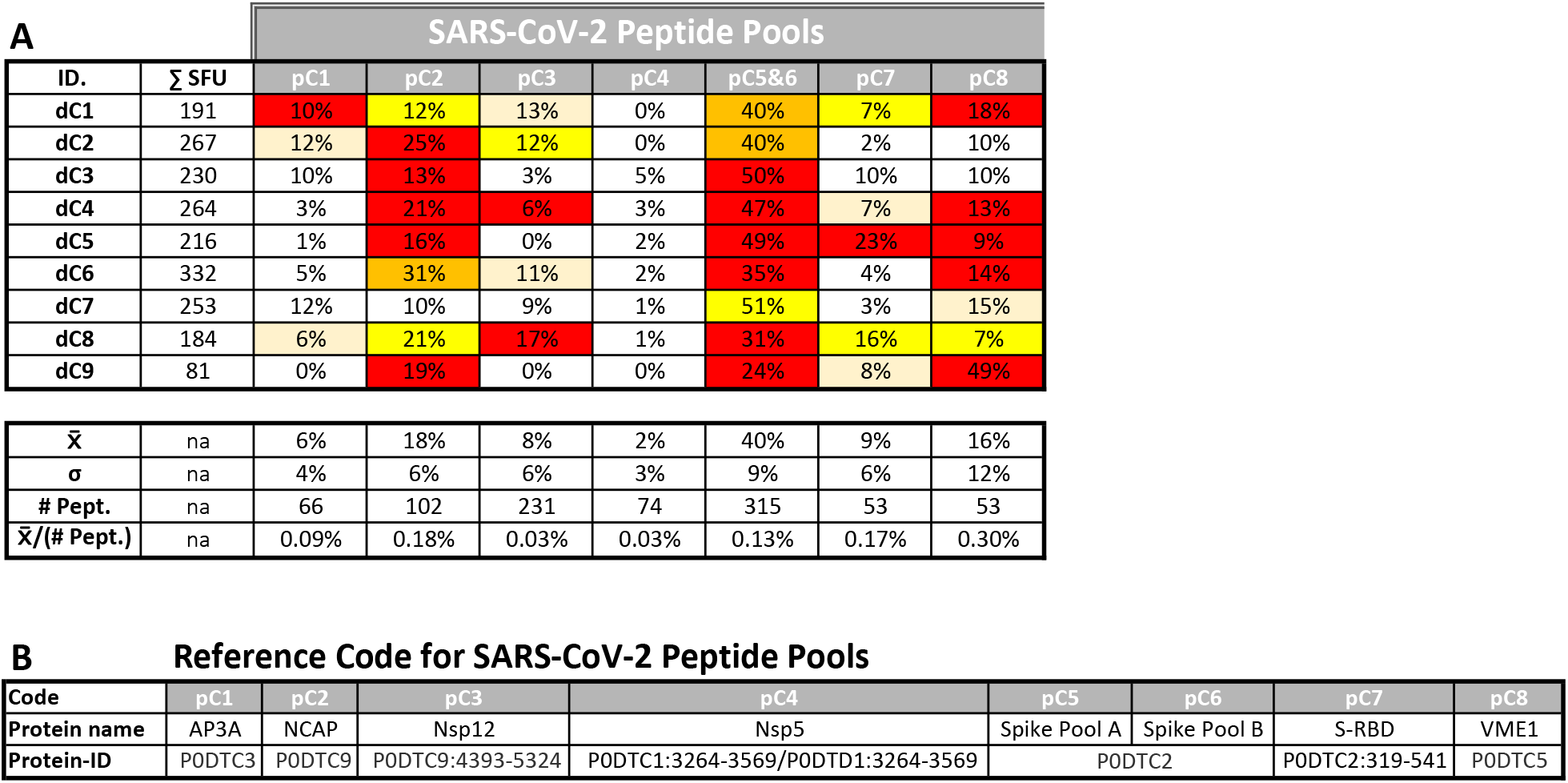
T cell immune dominance of SARS-CoV-2 proteins. The total SARS-CoV-2-specific T cell mass (ΣSFU) was calculated by adding up for each SARS-CoV-2-recovered donor the numbers of SFU elicited by all SARS-CoV-2 peptide pools in that donor at 1.5 μg/mL (see the raw data in Table 2). In the top panel, the percentage of T cells targeting each of the SARS-CoV-2 antigens is shown relative to the total clonal SARS-CoV-2-specific T cell mass in that individual, representing an immune dominance index. The superimposed heatmap specifies the affinity level of the respective T cell population, with the color code defined in Table 2. The lower panel shows the mean percentage 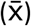 and SD of this immunedominance index for the cohort. Addressing the hypothesis that T cell immune dominance of a SARS-CoV-2 antigen is related to its size, the number of peptides in each pool is shown (# Pept.) and the mean immune dominance index 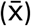 is normalized for the number of peptides 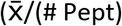.

One possible explanation for the relative immune dominance of Spike protein over the other SARS-CoV-2 proteins is its size relative to the others. The longer a protein, the more potential T cell epitopes it contains. Spike protein was covered by 315 peptides vs. for example NCAP, one third as long, which was covered by 102 peptides and VMA1, half as long as NCAP, with 53 peptides. Indeed, for these three antigens, and also for S-RBD and AP3A, the percentage of T cells targeting them divided by the number of peptides present in each pool (corresponding to the length of the respective protein) gave numbers in the same ballpark: 0.13%, 0.18%, 0.3%, 0.17%, and 0.09% for Spike, NCAP, VMA1, SRBD, and AP3A, respectively (Table 4). For these SARS-CoV-2 antigens, therefore, the magnitude of T cell response targeting each appeared to be a mere function of the proteins’ respective sizes. With this ratio substantially lower, at 0.03%, NSP12 and NSP5 were under-targeted relative to their size, possibly suggesting that the expression levels of these two antigens is lower during SARS-CoV-2 replication than that of the other SARS-CoV-2 antigens.

### 3.8. Non-Cross-Reactive T Cell Recognition of Seasonal Coronavirus Spike proteins

People around the world commonly get infected with seasonal coronaviruses such as 229E, NL63, OC43, and HKU1, and over the years most adults can be expected to have been infected with several of these coronavirus strains. T cell reactivity induced by SARS-CoV-2 mega peptide pools in individuals who clearly have not been exposed to SARS-CoV-2 have therefore been attributed to cognate cross-reactivity with seasonal coronavirus. By introducing negative control mega peptide pools to account for noise created by chance cross-reactivity (Table 1), and by adding the requirement for high affinity T cell recognition (Table 2), we show that SARS-CoV-2 antigens are not recognized by subjects in the Pre-COVID cohort as a consequence of cross-reactivity with seasonal coronavirus antigens. We therefore asked the reverse question: do mega peptide pools that specifically cover seasonal coronaviruses detect T cell memory in both cohorts?

We tested Spike proteins of 229E, NL63, OC43, and HKU1, which, due to their size, were each represented in two mega peptide pools (as was the SARS-CoV-2 Spike protein itself). These peptide pools were also tested at four concentrations, following exactly the same protocol as specified above for the SARS-CoV-2 Spike protein, and all other peptide pools tested in this study. While no Level 4 affinity response to SARS-CoV-2 Spike protein peptides were seen in Pre-COVID-19 subjects (Table 2B), eight of 20 subjects in this cohort exhibited a Level 4 response to at least one of the seasonal coronavirus Spike proteins (Table 5B). Cognate T cell reactivity to Spike proteins of seasonal coronaviruses, which do not cross-react with Spike protein of SARS-CoV-2, can therefore be detected in subjects bled in the Pre-COVID Era.

**Table 5.**
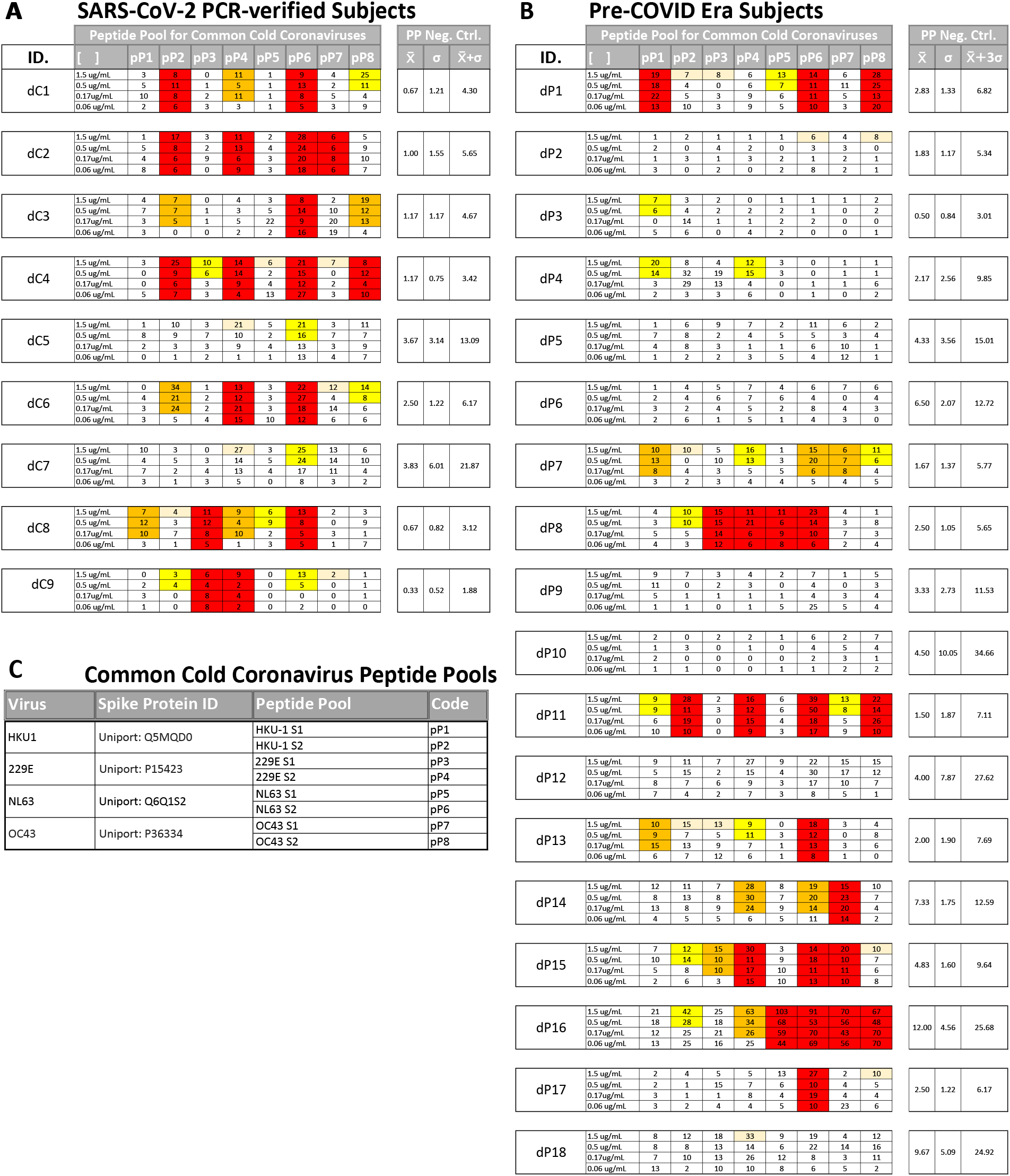
T cell recall responses to Spike proteins of the four Common Cold Coronaviruses, 229E, NL63, OC43, and HKU1, each represented due to size in two peptide pools. The code for the peptides is deciphered in S. Table 2. Each peptide pools has been tested in the specified four concentrations in a standard IFN-γ ELISPOT assay. SFU counts exceeding 3SD of the mean of the negative peptide pool control (PP. Neg. Contr.) shown on the left for each subject, are highlighted according to T cell affinity levels, as specified in Table 2.

Individuals in the SARS-CoV-2 PCR-verified cohort also showed Level 4 and Level 3 recall responses to Spike proteins of seasonal coronaviruses (Table 5A), however, with SFU numbers that were comparable with the Pre-COVID cohort. If these seasonal coronavirus-specific T cells cross-reacted with SARS-CoV-2 Spike protein, then they should occur in substantially elevated numbers compared to the Pre-COVID era subjects since the recent SARS-CoV-2 infection should have boosted such pre-existing memory cells and caused their numbers to selectively expand. Since seasonal coronavirus-reactive T cells occurred in comparable frequencies in both donor cohorts evaluated in this study, this further supports the data presented in Table 2 that peptides covering the SARS-CoV-2 Spike and seasonal coronavirus Spike antigens do not activate T cells in a cross-reactive fashion. Instead, T cells specifically recognizing the different coronavirus Spike proteins can be attributed to sequence disparities in the amino acid sequence of the Spike proteins expressed by these seasonal coronaviruses and the novel SARS-CoV-2 virus ^54, 65, 78^.

## 4. Concluding Remarks

The overall question that we addressed was whether test conditions can be established that permit to clearly identify SARS-CoV-2-specific T cell memory engaged in individuals who underwent a mild infection vs. humans who have not been infected with this virus. Previous publications on this subject matter reported up to 80% false positive results for uninfected individuals due to alleged T cell cross-reactivity. Here we have established criteria by which false positive results can be reduced to 0% (0 of 18 Pre-COVID Era test subjects), while permitting the detection of SARS-CoV-2-reactive T cells in eight of nine (89%) SARS-CoV-2 PCR-verified subjects. To accomplish this discrimination, a combination of four criteria needed to be used. First, the detection of ex vivo IFN-γ producing effector memory T cells. Second, we introduced negative control mega peptide pools, instead of medium alone, to establish the background noise level caused by chance cross-reactivity. Third, we introduced the criterion that a T cell response scores positive only if it has sufficient affinity, being triggered by at least four 1+2 (vol + vol) (3-fold) serial dilutions of the test peptides. Lastly, as COVID-recovered donors responded to several SARS-CoV-2 antigens, a broad T cell response profile was established as a requirement for scoring a subject positive. In addition to exhibiting a high affinity T cell response to at least one mega peptide pool, a second high or intermediate affinity level T cell response was also identified in 89% our SARS-CoV-2 PCR-verified cohort.

Following infection with the original SARS coronavirus (SARS-CoV), antibody and B cell memory wanes, but evidence for T cell memory remains ^79^. Antibody titers, and potentially B cell memory, also appear to be short-lived after SARS-CoV-2 infection as well ^80^, but it is presently not known whether T cell memory to this virus will be durably maintained. If serum antibody reactivity fails to provide reliable information on previous exposure to SARS-CoV-2, then potentially T cell diagnostics could fill this gap.

Even though we report here that SARS-CoV-2 and EBV-specific T cells occur in similar frequencies in COVID-recovered subjects (See Table 2 vs. S. Table 4), it might be premature to conclude that such findings signify the induction of a robust cognate T cell response following SARS-CoV-2 infection. Cognate T cell responses in general show a typical kinetic: in the first weeks after the onset of infection the frequency of the antigen-specific T cells reaches a peak, after which the frequencies drop to a substantially lower steady state level ^80^. We measured frequencies of SARS-CoV-2 antigen-specific T cells close to their expected peak in our COVID-recovered cohort, while the frequencies of the EBV-specific T cells were assessed in steady state. Therefore, in light of the typical T cell response kinetic, the data reported here, and supported by existing literature, may also signify that mild/asymptomatic SARS-CoV-2 infection induces a much weaker T cell response than natural EBV infection, or other viruses against which we develop protective immunity. The already low numbers of SARS-CoV-2-specific T cells early on after a mild/asymptomatic infection might further decrease with time, which needs to be established. Being able to accurately detect such rare SARS-CoV-2-specific T cells is an important step for immune diagnostics, but is just the first step toward understanding their role in host defense.

## Acknowledgements

We thank Ruliang Li of Cellular Technology Limited for expert technical assistance, Drs. Magdalena Tary-Lehmann, Nicholas Tomko and Alexey Y. Karulin for valuable discussions, and Diana Roen for editorial assistance. We also thank Melisa Sebok, Malachi Wickman, and Jennifer Penfold from American Red Cross, as well as Tibor Baki and Victoria Gaidaenko of CTL for helping us access blood from COVID-19-recovered subjects.

## Data Availability Statement

The datasets generated in this study will be made available by the authors, without undue reservation, to any qualified researcher.

## Author Contributions

Experiments were designed by A.A.L, PVL and P.A.R. Experimental data were generated by A.A.L, T.Z, and G.A.K.

