## Supplemental Figures for "Deconvoluting the T cell response to SARS-CoV-2: specificity versus chance- and cognate cross-reactivity"

**A** **SARS-CoV-2 PCR-verified Subjects**

| ID | PCR Confrimed | Hospitalization Yes or No | Days From Verification to Collect | DOC | Race | Gender | Age |
|---|---|---|---|---|---|---|---|
| dC1 | Pos | No | 14 | 05/11/20 | African/American | Male | 49 |
| dC2 | Pos | No | 14 | 05/11/20 | African/American | Female | 20 |
| dC3 | Pos | No | 34 | 06/11/20 | Caucasian | Female | 28 |
| dC4 | Pos | No | 34 | 7/7/2020 | Caucasian | Female | 24 |
| dC5 | Pos | No | 24 | 7/20/2020 | African/American | Male | 33 |
| dC6 | Pos | No | 20 | 7/22/2020 | Hispanic/Latino | Female | 25 |
| dC7 | Pos | No | 17 | 05/14/20 | African/American | Female | 22 |
| dC8 | Pos | Yes | 31 | 7/6/2020 | Caucasian | Female | 53 |
| dC9 | Pos | No | 95 | 07/08/20 | Caucasian | Female | 51 |

**B** **Pre-COVID Era Subjects**

| ID | DOC | Race | Gender | Age |
|---|---|---|---|---|
| dP1 | 3/15/2017 | Hispanic | Male | 18 |
| dP2 | 9/27/2017 | African/American | Female | 28 |
| dP3 | 7/26/2017 | Hispanic | Male | 34 |
| dP4 | 5/16/2018 | Hispanic | Male | 37 |
| dP5 | 10/16/2018 | Hispanic | Male | 30 |
| dP6 | 10/1/2018 | Caucasian | Female | 34 |
| dP7 | 3/14/2018 | Hispanic | Male | 31 |
| dP8 | 8/16/2017 | Hispanic | Female | 39 |
| dP9 | 8/13/2018 | African/American | Male | 49 |
| dP10 | 7/20/2016 | Caucasian | Female | 43 |
| dP11 | 8/22/2019 | Caucasian | Female | 22 |
| dP12 | 7/15/2019 | Hispanic | Male | 24 |
| dP13 | 6/12/2019 | Hispanic | Female | 52 |
| dP14 | 5/6/2019 | Hispanic/Latino | Female | 46 |
| dP15 | 4/10/2019 | Hispanic/Latino | Male | 36 |
| dP16 | 11/2/2011 | Hispanic | Male | 42 |
| dP17 | 7/8/2019 | Caucasian | Male | 54 |
| dP18 | 7/16/2017 | Hispanic | Male | 45 |

S. Table 1. Human subjects tested in this study. The Pre-COVID Era cohort (B) consisted of 18 subjects who were bled prior to December 2019 (subject IDs **dP**1-18, for **d**onor **P**re-COVID-19 Era). The SARS-CoV-2-PCR-verified cohort (A) consisted of nine individuals who underwent mild PCR-confirmed SARS-CoV-2 infection (subject IDs **dC**1-9, for **d**onors recovered from **C**OVID-19).

| Peptide Pools for SARS-CoV-2 Proteins | | | | | |
|---|---|---|---|---|---|
| **Code** | **Protein name** | **Origin** | **Protein-ID** | **# of Peptides** | **Product Code** |
| pC1 | AP3A | SAR-CoV-2 | P0DTC3 | 66 | PM-WCPV-AP3A |
| pC2 | NCAP | | P0DTC9 | 102 | PM-WCPV-NCAP |
| pC3 | Nsp12 | | P0DTC9:4393-5324 | 231 | PM-WCPV-Nsp12-1 |
| pC4 | Nsp5 | | P0DTC1:3264-3569 | 74 | PM-WCPV-Nsp5-1 |
| pC5 | Spike Pool A | | P0DTC2 | 158 | PM-WCPV-S |
| pC6 | Spile Pool B | | | 157 | |
| pC7 | S-RBD | | P0DTC2:319-541 | 53 | PM-WCPV-S-RBD-1 |
| pC8 | VME1 | | P0DTC5 | 53 | PM-WCPV-VME |

| Peptide Pools for EBV Proteins | | | | | |
|---|---|---|---|---|---|
| **Code** | **Protein name** | **Origin** | **Protein-ID** | **# of Peptides** | **Product Code** |
| pE1 | BARF1 | EBV | P03228 | 53 | PM-C-EBV-2 |
| pE2 | BMLF1 | | Q04360 | 117 | |
| pE3 | BMRF1 | | P03191 | 99 | |
| pE4 | BRLF1 | | P03209 | 149 | |
| pE5 | BZLF1 | | P03206 | 59 | |
| pE6 | EBNA-LP | | Q8AZK7 | 124 | |
| pE7 | EBNA1 | | P03211 | 158 | |
| pE8 | EBNA2 | | P12978 | 19 | |
| pE9 | EBNA3a | | P12977 | 234 | |
| pE10 | EBNA3b | | Q1HVG4 | 279 | |
| pE11 | EBNA3c | | Q69140 | 265 | |
| pE12 | GP350/340 | | P03200 | 224 | |
| pE13 | LMP1 | | PO3230 | 94 | |
| pE14 | LMP2 | | P13285 | 122 | |

| Candidate Negative Control Peptide Pools | | | | | |
|---|---|---|---|---|---|
| **Code** | **Protein name** | **Origin** | **Protein-ID** | **# of Peptides** | **Product Code** |
| pN1 | Nef Protein | HIV | LAN* | 150 | PM-HIV-NEF |
| pN2 | Gag polyprotein | HIV | LAN* | 150 | PM-HIV-GAG |
| pN3 | Pol Polyprotein | HIV | LAN* | 150 | PM-HIV-POL |
| pN4 | Con B gag motif | HIV | LAN* | 123 | PM-HIV-CONB |
| pN5 | Actin | Human | P68133** | 92 | PM-ACTS |
| pN6 | Nucleoprotein | Ebola-Thai | B8XCN6** | 182 | PM-TEBOV-NP |
| pN7 | ENV GP | HIV | LAN* | 150 | PM-HIV-ENV |

| Peptide Pools for S Proteins of Common Cold Coronaviruses | | | | | |
|---|---|---|---|---|---|
| **Origin** | **Protein-ID** | **Product Code** | **Peptide Pool** | **# of Peptides** | **Code** |
| HKU1 | Q5MQD0 | PM-HKU1-S-1 | HKU-1 S1 | 169 | pP1 |
| | | | HKU-1 S2 | 168 | pP2 |
| HCoV-229E | P15423 | PM-229E-S-1 | 229E S1 | 146 | pP3 |
| | | | 229E S2 | 145 | pP4 |
| HCoV-NL63 | Q6Q1S2 | PM-NL63-S-1 | NL63 S1 | 169 | pP5 |
| | | | NL63 S2 | 168 | pP6 |
| HCoV-OC43 | P36334 | PM-OC43-S-1 | OC43 S1 | 168 | pP7 |
| | | | OC43 S2 | 168 | pP8 |

S. Table 2. Definition of mega peptide pools used in this study. All peptide pools consisted of unpurified 15-mer peptides that systematically cover the entire amino acid (a.a) sequence of the respective protein in steps of 11 a.a. The number of peptides in each pool is specified. All of these peptide pools are commercially available from JPT.

**A** Positive Responses Based on Media Control Background

| ID. | [ ] | pC1 | pC2 | pC3 | pC4 | pC4 | pC5 | pC6 | pC7 | Media | $\bar{x}$ | σ | $\bar{x}+3\sigma$ |
|---|---|---|---|---|---|---|---|---|---|---|---|---|---|
| | | SARS-CoV-2 Peptide Pools | | | | | | | | Media Controls | | | |
| dP1 | 1.5 ug/mL | 0 | 5 | 2 | 3 | 5 | 8 | 4 | 2 | 1 | 1.25 | 0.50 | 2.75 |
| | 0.5 ug/mL | 0 | 3 | 3 | 1 | 6 | 7 | 8 | 1 | 2 | | | |
| | 0.17ug/mL | 0 | 3 | 4 | 0 | 3 | 7 | 0 | 2 | 1 | | | |
| | 0.06 ug/mL | 1 | 5 | 3 | 6 | 2 | 5 | 5 | 1 | 1 | | | |
| dP2 | 1.5 ug/mL | 0 | 0 | 3 | 3 | 1 | 6 | 2 | 1 | 0 | 0.50 | 1.00 | 3.50 |
| | 0.5 ug/mL | 2 | 2 | 1 | 2 | 3 | 0 | 4 | 1 | 0 | | | |
| | 0.17ug/mL | 4 | 2 | 2 | 1 | 2 | 3 | 2 | 3 | 2 | | | |
| | 0.06 ug/mL | 2 | 3 | 2 | 1 | 0 | 2 | 1 | 3 | 0 | | | |
| dP3 | 1.5 ug/mL | 0 | 4 | 7 | 1 | 1 | 6 | 3 | 1 | 0 | 1.50 | 1.73 | 6.70 |
| | 0.5 ug/mL | 2 | 2 | 10 | 0 | 4 | 4 | 1 | 3 | 3 | | | |
| | 0.17ug/mL | 1 | 1 | 4 | 1 | 5 | 2 | 1 | 1 | 3 | | | |
| | 0.06 ug/mL | 4 | 1 | 3 | 0 | 1 | 3 | 1 | 1 | 0 | | | |
| dP4 | 1.5 ug/mL | 12 | 6 | 11 | 5 | 8 | 35 | 6 | 3 | 0 | 2.50 | 2.89 | 11.16 |
| | 0.5 ug/mL | 15 | 14 | 11 | 7 | 7 | 28 | 0 | 13 | 5 | | | |
| | 0.17ug/mL | 4 | 7 | 6 | 10 | 4 | 7 | 1 | 8 | 5 | | | |
| | 0.06 ug/mL | 6 | 12 | 5 | 1 | 4 | 4 | 2 | 1 | 0 | | | |
| dP5 | 1.5 ug/mL | 5 | 1 | 3 | 2 | 1 | 1 | 0 | 0 | 3 | 1.25 | 1.26 | 5.02 |
| | 0.5 ug/mL | 2 | 0 | 0 | 2 | 1 | 3 | 1 | 4 | 1 | | | |
| | 0.17ug/mL | 1 | 1 | 2 | 3 | 0 | 1 | 1 | 3 | 0 | | | |
| | 0.06 ug/mL | 2 | 2 | 2 | 1 | 2 | 0 | 2 | 2 | 1 | | | |
| dP6 | 1.5 ug/mL | 2 | 3 | 4 | 4 | 3 | 1 | 4 | 1 | 4 | 1.75 | 1.71 | 6.87 |
| | 0.5 ug/mL | 2 | 4 | 5 | 6 | 3 | 4 | 8 | 2 | 1 | | | |
| | 0.17ug/mL | 4 | 5 | 2 | 3 | 0 | 4 | 1 | 6 | 2 | | | |
| | 0.06 ug/mL | 0 | 3 | 2 | 6 | 6 | 3 | 3 | 2 | 0 | | | |
| dP7 | 1.5 ug/mL | 3 | 0 | 4 | 1 | 2 | 2 | 4 | 3 | 1 | 0.75 | 0.50 | 2.25 |
| | 0.5 ug/mL | 5 | 1 | 3 | 2 | 4 | 5 | 1 | 4 | 1 | | | |
| | 0.17ug/mL | 1 | 0 | 0 | 2 | 1 | 3 | 1 | 1 | 0 | | | |
| | 0.06 ug/mL | 1 | 1 | 1 | 0 | 1 | 1 | 2 | 1 | 1 | | | |
| dP8 | 1.5 ug/mL | 6 | 0 | 1 | 0 | 5 | 4 | 1 | 4 | 1 | 2.50 | 3.11 | 11.83 |
| | 0.5 ug/mL | 3 | 1 | 1 | 3 | 5 | 3 | 4 | 5 | 2 | | | |
| | 0.17ug/mL | 7 | 0 | 2 | 1 | 10 | 4 | 4 | 4 | 0 | | | |
| | 0.06 ug/mL | 5 | 0 | 1 | 1 | 3 | 3 | 3 | 4 | 7 | | | |
| dP9 | 1.5 ug/mL | 1 | 0 | 1 | 0 | 0 | 0 | 2 | 2 | 2 | 0.50 | 1.00 | 3.50 |
| | 0.5 ug/mL | 0 | 0 | 1 | 0 | 0 | 1 | 0 | 0 | 0 | | | |
| | 0.17ug/mL | 0 | 3 | 0 | 0 | 0 | 1 | 1 | 0 | 0 | | | |
| | 0.06 ug/mL | 1 | 0 | 0 | 0 | 0 | 3 | 1 | 1 | 0 | | | |
| dP10 | 1.5 ug/mL | 0 | 0 | 0 | 0 | 0 | 1 | 0 | 0 | 0 | 0.25 | 0.50 | 1.75 |
| | 0.5 ug/mL | 0 | 0 | 0 | 0 | 2 | 2 | 0 | 0 | 1 | | | |
| | 0.17ug/mL | 0 | 0 | 0 | 0 | 0 | 0 | 0 | 0 | 0 | | | |
| | 0.06 ug/mL | 1 | 1 | 0 | 2 | 0 | 1 | 0 | 0 | 0 | | | |
| dP11 | 1.5 ug/mL | 1 | 3 | 2 | 0 | 5 | 4 | 4 | 1 | 1 | 0.50 | 0.58 | 2.23 |
| | 0.5 ug/mL | 0 | 3 | 1 | 3 | 1 | 2 | 0 | 0 | 0 | | | |
| | 0.17ug/mL | 4 | 1 | 3 | 0 | 2 | 3 | 0 | 0 | 0 | | | |
| | 0.06 ug/mL | 1 | 2 | 2 | 1 | 0 | 3 | 0 | 0 | 1 | | | |
| dP12 | 1.5 ug/mL | 0 | 1 | 2 | 3 | 4 | 3 | 3 | 1 | 1 | 1.00 | 0.82 | 3.45 |
| | 0.5 ug/mL | 2 | 1 | 4 | 4 | 2 | 5 | 1 | 0 | 1 | | | |
| | 0.17ug/mL | 1 | 0 | 2 | 0 | 0 | 1 | 0 | 2 | 0 | | | |
| | 0.06 ug/mL | 1 | 0 | 0 | 0 | 7 | 4 | 1 | 2 | 2 | | | |
| dP13 | 1.5 ug/mL | 0 | 1 | 2 | 1 | 0 | 0 | 0 | 0 | 0 | 1.00 | 1.41 | 5.24 |
| | 0.5 ug/mL | 2 | 1 | 0 | 2 | 2 | 0 | 1 | 1 | 0 | | | |
| | 0.17ug/mL | 0 | 0 | 0 | 0 | 1 | 1 | 6 | 1 | 1 | | | |
| | 0.06 ug/mL | 0 | 1 | 3 | 0 | 1 | 0 | 1 | 3 | 3 | | | |
| dP14 | 1.5 ug/mL | 1 | 6 | 2 | 8 | 1 | 3 | 2 | 4 | 2 | 4.25 | 3.30 | 14.16 |
| | 0.5 ug/mL | 4 | 6 | 2 | 2 | 3 | 5 | 3 | 7 | 8 | | | |
| | 0.17ug/mL | 3 | 6 | 3 | 5 | 8 | 3 | 2 | 4 | 1 | | | |
| | 0.06 ug/mL | 2 | 6 | 3 | 1 | 4 | 1 | 1 | 1 | 6 | | | |
| dP15 | 1.5 ug/mL | 7 | 5 | 6 | 4 | 7 | 10 | 6 | 5 | 5 | 4.75 | 2.87 | 13.37 |
| | 0.5 ug/mL | 5 | 6 | 1 | 3 | 3 | 2 | 7 | 2 | 1 | | | |
| | 0.17ug/mL | 3 | 4 | 8 | 5 | 5 | 7 | 4 | 0 | 5 | | | |
| | 0.06 ug/mL | 4 | 7 | 3 | 3 | 7 | 7 | 9 | 6 | 8 | | | |
| dP16 | 1.5 ug/mL | 13 | 18 | 22 | 7 | 23 | 16 | 15 | 11 | 18 | 15.67 | 4.04 | 27.79 |
| | 0.5 ug/mL | 8 | 11 | 13 | 11 | 20 | 30 | 11 | 3 | 18 | | | |
| | 0.17ug/mL | 7 | 6 | 8 | 7 | 14 | 9 | 9 | 2 | 11 | | | |
| | 0.06 ug/mL | 10 | 11 | 11 | 12 | 15 | 15 | 16 | 7 | 56 | | | |
| dP17 | 1.5 ug/mL | 0 | 0 | 1 | 0 | 0 | 3 | 0 | 4 | 0 | 3.50 | 3.87 | 15.12 |
| | 0.5 ug/mL | 2 | 0 | 4 | 2 | 1 | 2 | 1 | 1 | 3 | | | |
| | 0.17ug/mL | 0 | 1 | 0 | 3 | 0 | 2 | 1 | 7 | 2 | | | |
| | 0.06 ug/mL | 1 | 2 | 1 | 3 | 0 | 2 | 2 | 4 | 9 | | | |
| dP18 | 1.5 ug/mL | 0 | 4 | 1 | 2 | 9 | 6 | 5 | 6 | 2 | 2.25 | 1.71 | 7.37 |
| | 0.5 ug/mL | 0 | 2 | 1 | 2 | 9 | 9 | 7 | 10 | 4 | | | |
| | 0.17ug/mL | 0 | 4 | 4 | 5 | 9 | 7 | 10 | 5 | 0 | | | |
| | 0.06 ug/mL | 0 | 1 | 4 | 4 | 3 | 8 | 3 | 5 | 3 | | | |
| dC1 | 1.5 ug/mL | 12 | 15 | 16 | 0 | 27 | 22 | 9 | 22 | 8 | 2.00 | 4.00 | 14.00 |
| | 0.5 ug/mL | 13 | 6 | 2 | 0 | 22 | 5 | 5 | 19 | 0 | | | |
| | 0.17ug/mL | 17 | 4 | 0 | 0 | 6 | 9 | 2 | 27 | 0 | | | |
| | 0.06 ug/mL | 13 | 2 | 2 | 0 | 3 | 4 | 2 | 26 | 0 | | | |
| dC2 | 1.5 ug/mL | 6 | 13 | 6 | 0 | 6 | 15 | 1 | 5 | 1 | 1.00 | 0.82 | 3.45 |
| | 0.5 ug/mL | 2 | 16 | 6 | 2 | 9 | 8 | 1 | 5 | 0 | | | |
| | 0.17ug/mL | 3 | 16 | 1 | 0 | 5 | 7 | 1 | 1 | 2 | | | |
| | 0.06 ug/mL | 1 | 6 | 3 | 1 | 6 | 4 | 0 | 7 | 1 | | | |
| dC3 | 1.5 ug/mL | 4 | 5 | 1 | 2 | 15 | 5 | 4 | 4 | 0 | 0.25 | 0.50 | 1.75 |
| | 0.5 ug/mL | 3 | 7 | 0 | 4 | 22 | 11 | 9 | 1 | 1 | | | |
| | 0.17ug/mL | 3 | 6 | 1 | 1 | 16 | 5 | 2 | 2 | 0 | | | |
| | 0.06 ug/mL | 1 | 5 | 1 | 1 | 11 | 10 | 4 | 3 | 0 | | | |
| dC4 | 1.5 ug/mL | 2 | 14 | 4 | 2 | 7 | 25 | 5 | 9 | 2 | 0.75 | 0.96 | 3.62 |
| | 0.5 ug/mL | 3 | 18 | 5 | 3 | 8 | 33 | 3 | 11 | 1 | | | |
| | 0.17ug/mL | 3 | 6 | 6 | 3 | 8 | 24 | 8 | 6 | 0 | | | |
| | 0.06 ug/mL | 2 | 6 | 38 | 1 | 4 | 15 | 3 | 13 | 0 | | | |
| dC5 | 1.5 ug/mL | 5 | 61 | 0 | 6 | 143 | 45 | 90 | 34 | 4 | 1.50 | 1.91 | 7.24 |
| | 0.5 ug/mL | 0 | 53 | 3 | 5 | 105 | 40 | 60 | 41 | 0 | | | |
| | 0.17ug/mL | 5 | 43 | 8 | 6 | 85 | 24 | 26 | 35 | 0 | | | |
| | 0.06 ug/mL | 5 | 48 | 4 | 4 | 40 | 19 | 22 | 30 | 2 | | | |
| dC6 | 1.5 ug/mL | 5 | 34 | 12 | 2 | 18 | 20 | 4 | 15 | 0 | 1.75 | 1.26 | 5.52 |
| | 0.5 ug/mL | 3 | 23 | 2 | 4 | 31 | 19 | 2 | 14 | 3 | | | |
| | 0.17ug/mL | 7 | 15 | 2 | 2 | 14 | 19 | 1 | 11 | 2 | | | |
| | 0.06 ug/mL | 1 | 4 | 6 | 5 | 15 | 19 | 5 | 15 | 2 | | | |
| dC7 | 1.5 ug/mL | 17 | 14 | 13 | 1 | 30 | 45 | 4 | 22 | 0 | 1.25 | 1.26 | 5.02 |
| | 0.5 ug/mL | 14 | 12 | 4 | 0 | 23 | 28 | 7 | 17 | 1 | | | |
| | 0.17ug/mL | 6 | 7 | 0 | 0 | 5 | 10 | 4 | 17 | 1 | | | |
| | 0.06 ug/mL | 7 | 6 | 0 | 2 | 5 | 4 | 5 | 11 | 3 | | | |
| dC8 | 1.5 ug/mL | 4 | 15 | 12 | 1 | 8 | 14 | 11 | 5 | 0 | 1.00 | 1.41 | 5.24 |
| | 0.5 ug/mL | 0 | 4 | 16 | 2 | 8 | 18 | 2 | 5 | 1 | | | |
| | 0.17ug/mL | 1 | 2 | 9 | 0 | 4 | 19 | 3 | 1 | 3 | | | |
| | 0.06 ug/mL | 1 | 5 | 7 | 0 | 4 | 11 | 8 | 2 | 0 | | | |
| dC9 | 1.5 ug/mL | 0 | 17 | 0 | 0 | 17 | 5 | 7 | 44 | 0 | 0.00 | 0.00 | 0.00 |
| | 0.5 ug/mL | 4 | 12 | 0 | 0 | 12 | 9 | 1 | 36 | 0 | | | |
| | 0.17ug/mL | 4 | 9 | 1 | 0 | 6 | 4 | 1 | 17 | 0 | | | |
| | 0.06 ug/mL | 2 | 8 | 0 | 0 | 7 | 1 | 3 | 12 | 0 | | | |

**B** Positive Responsese Based on Negative Control Peptide Background

| ID. | [ ] | pC1 | pC2 | pC3 | pC4 | pC4 | pC5 | pC6 | pC7 | $\bar{x}$ | σ | $\bar{x}+3\sigma$ |
|---|---|---|---|---|---|---|---|---|---|---|---|---|
| | | SARS-CoV-2 Peptide Pools | | | | | | | | PP Neg. Ctrl. | | |
| dP1 | 1.5 ug/mL | 0 | 5 | 2 | 3 | 5 | 8 | 4 | 2 | 2.83 | 1.33 | 6.82 |
| | 0.5 ug/mL | 0 | 3 | 3 | 1 | 6 | 7 | 8 | 1 | | | |
| | 0.17ug/mL | 0 | 3 | 4 | 0 | 3 | 7 | 0 | 2 | | | |
| | 0.06 ug/mL | 1 | 5 | 3 | 6 | 2 | 5 | 5 | 1 | | | |
| dP2 | 1.5 ug/mL | 0 | 0 | 3 | 3 | 1 | 6 | 2 | 1 | 1.83 | 1.17 | 5.34 |
| | 0.5 ug/mL | 2 | 2 | 1 | 2 | 3 | 0 | 4 | 1 | | | |
| | 0.17ug/mL | 4 | 2 | 2 | 1 | 2 | 3 | 2 | 3 | | | |
| | 0.06 ug/mL | 2 | 3 | 2 | 1 | 0 | 2 | 1 | 3 | | | |
| dP3 | 1.5 ug/mL | 0 | 4 | 7 | 1 | 1 | 6 | 3 | 1 | 0.50 | 0.84 | 3.01 |
| | 0.5 ug/mL | 2 | 2 | 10 | 0 | 4 | 4 | 1 | 3 | | | |
| | 0.17ug/mL | 1 | 1 | 4 | 1 | 5 | 2 | 1 | 1 | | | |
| | 0.06 ug/mL | 4 | 1 | 3 | 0 | 1 | 3 | 1 | 1 | | | |
| dP4 | 1.5 ug/mL | 12 | 6 | 11 | 5 | 8 | 35 | 6 | 3 | 2.17 | 2.56 | 9.85 |
| | 0.5 ug/mL | 15 | 14 | 11 | 7 | 7 | 28 | 0 | 13 | | | |
| | 0.17ug/mL | 4 | 7 | 6 | 10 | 4 | 7 | 1 | 8 | | | |
| | 0.06 ug/mL | 6 | 12 | 5 | 1 | 4 | 4 | 2 | 1 | | | |
| dP5 | 1.5 ug/mL | 5 | 1 | 3 | 2 | 1 | 1 | 0 | 0 | 4.33 | 3.56 | 15.01 |
| | 0.5 ug/mL | 2 | 0 | 0 | 2 | 1 | 3 | 1 | 4 | | | |
| | 0.17ug/mL | 1 | 1 | 2 | 3 | 0 | 1 | 1 | 3 | | | |
| | 0.06 ug/mL | 2 | 2 | 2 | 1 | 2 | 0 | 2 | 2 | | | |
| dP6 | 1.5 ug/mL | 2 | 3 | 4 | 4 | 3 | 1 | 4 | 1 | 6.50 | 2.07 | 12.72 |
| | 0.5 ug/mL | 2 | 4 | 5 | 6 | 3 | 4 | 8 | 2 | | | |
| | 0.17ug/mL | 4 | 5 | 2 | 3 | 0 | 4 | 1 | 6 | | | |
| | 0.06 ug/mL | 0 | 3 | 2 | 6 | 6 | 3 | 3 | 2 | | | |
| dP7 | 1.5 ug/mL | 3 | 0 | 4 | 1 | 2 | 2 | 4 | 3 | 1.67 | 1.37 | 5.77 |
| | 0.5 ug/mL | 5 | 1 | 3 | 2 | 4 | 5 | 1 | 4 | | | |
| | 0.17ug/mL | 1 | 0 | 0 | 2 | 1 | 3 | 1 | 1 | | | |
| | 0.06 ug/mL | 1 | 1 | 1 | 0 | 1 | 1 | 2 | 1 | | | |
| dP8 | 1.5 ug/mL | 6 | 0 | 1 | 0 | 5 | 4 | 1 | 4 | 2.50 | 1.05 | 5.65 |
| | 0.5 ug/mL | 3 | 1 | 1 | 3 | 5 | 3 | 4 | 5 | | | |
| | 0.17ug/mL | 7 | 0 | 2 | 1 | 10 | 4 | 4 | 4 | | | |
| | 0.06 ug/mL | 5 | 0 | 1 | 1 | 3 | 3 | 3 | 4 | | | |
| dP9 | 1.5 ug/mL | 1 | 0 | 1 | 0 | 0 | 0 | 2 | 2 | 3.33 | 2.73 | 11.53 |
| | 0.5 ug/mL | 0 | 0 | 1 | 0 | 0 | 1 | 0 | 0 | | | |
| | 0.17ug/mL | 0 | 3 | 0 | 0 | 0 | 1 | 1 | 0 | | | |
| | 0.06 ug/mL | 1 | 0 | 0 | 0 | 0 | 3 | 1 | 1 | | | |
| dP10 | 1.5 ug/mL | 0 | 0 | 0 | 0 | 0 | 1 | 0 | 0 | 4.50 | 10.05 | 34.66 |
| | 0.5 ug/mL | 0 | 0 | 0 | 0 | 2 | 2 | 0 | 0 | | | |
| | 0.17ug/mL | 0 | 0 | 0 | 0 | 0 | 0 | 0 | 0 | | | |
| | 0.06 ug/mL | 1 | 1 | 0 | 2 | 0 | 1 | 0 | 0 | | | |
| dP11 | 1.5 ug/mL | 1 | 3 | 2 | 0 | 5 | 4 | 4 | 1 | 1.50 | 1.87 | 7.11 |
| | 0.5 ug/mL | 0 | 3 | 1 | 3 | 1 | 2 | 0 | 0 | | | |
| | 0.17ug/mL | 4 | 1 | 3 | 0 | 2 | 3 | 0 | 0 | | | |
| | 0.06 ug/mL | 1 | 2 | 2 | 1 | 0 | 3 | 0 | 0 | | | |
| dP12 | 1.5 ug/mL | 0 | 1 | 2 | 3 | 4 | 3 | 3 | 1 | 4.00 | 7.87 | 27.62 |
| | 0.5 ug/mL | 2 | 1 | 4 | 4 | 2 | 5 | 1 | 0 | | | |
| | 0.17ug/mL | 1 | 0 | 2 | 0 | 0 | 1 | 0 | 2 | | | |
| | 0.06 ug/mL | 1 | 0 | 0 | 0 | 7 | 4 | 1 | 2 | | | |
| dP13 | 1.5 ug/mL | 0 | 1 | 2 | 1 | 0 | 0 | 0 | 0 | 2.00 | 1.90 | 7.69 |
| | 0.5 ug/mL | 2 | 1 | 0 | 2 | 2 | 0 | 1 | 1 | | | |
| | 0.17ug/mL | 0 | 0 | 0 | 0 | 1 | 1 | 6 | 1 | | | |
| | 0.06 ug/mL | 0 | 1 | 3 | 0 | 1 | 0 | 1 | 3 | | | |
| dP14 | 1.5 ug/mL | 1 | 6 | 2 | 8 | 1 | 3 | 2 | 4 | 7.33 | 1.75 | 12.59 |
| | 0.5 ug/mL | 4 | 6 | 2 | 2 | 3 | 5 | 3 | 7 | | | |
| | 0.17ug/mL | 3 | 6 | 3 | 5 | 8 | 3 | 2 | 4 | | | |
| | 0.06 ug/mL | 2 | 6 | 3 | 1 | 4 | 1 | 1 | 1 | | | |
| dP15 | 1.5 ug/mL | 7 | 5 | 6 | 4 | 7 | 10 | 6 | 5 | 4.83 | 1.60 | 9.64 |
| | 0.5 ug/mL | 5 | 6 | 1 | 3 | 3 | 2 | 7 | 2 | | | |
| | 0.17ug/mL | 3 | 4 | 8 | 5 | 5 | 7 | 4 | 0 | | | |
| | 0.06 ug/mL | 4 | 7 | 3 | 3 | 7 | 7 | 9 | 6 | | | |
| dP16 | 1.5 ug/mL | 13 | 18 | 22 | 7 | 23 | 16 | 15 | 11 | 12.00 | 4.56 | 25.68 |
| | 0.5 ug/mL | 8 | 11 | 13 | 11 | 20 | 30 | 11 | 3 | | | |
| | 0.17ug/mL | 7 | 6 | 8 | 7 | 14 | 9 | 9 | 2 | | | |
| | 0.06 ug/mL | 10 | 11 | 11 | 12 | 15 | 15 | 16 | 7 | | | |
| dP17 | 1.5 ug/mL | 0 | 0 | 1 | 0 | 0 | 3 | 0 | 4 | 2.50 | 1.22 | 6.17 |
| | 0.5 ug/mL | 2 | 0 | 4 | 2 | 1 | 2 | 1 | 1 | | | |
| | 0.17ug/mL | 0 | 1 | 0 | 3 | 0 | 2 | 1 | 7 | | | |
| | 0.06 ug/mL | 1 | 2 | 1 | 3 | 0 | 2 | 2 | 4 | | | |
| dP18 | 1.5 ug/mL | 0 | 4 | 1 | 2 | 9 | 6 | 5 | 6 | 9.67 | 5.09 | 24.92 |
| | 0.5 ug/mL | 0 | 2 | 1 | 2 | 9 | 9 | 7 | 10 | | | |
| | 0.17ug/mL | 0 | 4 | 4 | 5 | 9 | 7 | 10 | 5 | | | |
| | 0.06 ug/mL | 0 | 1 | 4 | 4 | 3 | 8 | 3 | 5 | | | |
| dC1 | 1.5 ug/mL | 12 | 15 | 16 | 0 | 27 | 22 | 9 | 22 | 0.67 | 1.21 | 4.30 |
| | 0.5 ug/mL | 13 | 6 | 2 | 0 | 22 | 5 | 5 | 19 | | | |
| | 0.17ug/mL | 17 | 4 | 0 | 0 | 6 | 9 | 2 | 27 | | | |
| | 0.06 ug/mL | 13 | 2 | 2 | 0 | 3 | 4 | 2 | 26 | | | |
| dC2 | 1.5 ug/mL | 6 | 13 | 6 | 0 | 6 | 15 | 1 | 5 | 1.00 | 1.55 | 5.65 |
| | 0.5 ug/mL | 2 | 16 | 6 | 2 | 9 | 8 | 1 | 5 | | | |
| | 0.17ug/mL | 3 | 16 | 1 | 0 | 5 | 7 | 1 | 1 | | | |
| | 0.06 ug/mL | 1 | 6 | 3 | 1 | 6 | 4 | 0 | 7 | | | |
| dC3 | 1.5 ug/mL | 4 | 5 | 1 | 2 | 15 | 5 | 4 | 4 | 1.17 | 1.17 | 4.67 |
| | 0.5 ug/mL | 3 | 7 | 0 | 4 | 22 | 11 | 9 | 1 | | | |
| | 0.17ug/mL | 3 | 6 | 1 | 1 | 16 | 5 | 2 | 2 | | | |
| | 0.06 ug/mL | 1 | 5 | 1 | 1 | 11 | 10 | 4 | 3 | | | |
| dC4 | 1.5 ug/mL | 2 | 14 | 4 | 2 | 7 | 25 | 5 | 9 | 1.17 | 0.75 | 3.42 |
| | 0.5 ug/mL | 3 | 18 | 5 | 3 | 8 | 33 | 3 | 11 | | | |
| | 0.17ug/mL | 3 | 6 | 6 | 3 | 8 | 24 | 8 | 6 | | | |
| | 0.06 ug/mL | 2 | 6 | 38 | 1 | 4 | 15 | 3 | 13 | | | |
| dC5 | 1.5 ug/mL | 5 | 61 | 0 | 6 | 143 | 45 | 90 | 34 | 3.67 | 3.14 | 13.09 |
| | 0.5 ug/mL | 0 | 53 | 3 | 5 | 105 | 40 | 60 | 41 | | | |
| | 0.17ug/mL | 5 | 43 | 8 | 6 | 85 | 24 | 26 | 35 | | | |
| | 0.06 ug/mL | 5 | 48 | 4 | 4 | 40 | 19 | 22 | 30 | | | |
| dC6 | 1.5 ug/mL | 5 | 34 | 12 | 2 | 18 | 20 | 4 | 15 | 2.50 | 1.22 | 6.17 |
| | 0.5 ug/mL | 3 | 23 | 2 | 4 | 31 | 19 | 2 | 14 | | | |
| | 0.17ug/mL | 7 | 15 | 2 | 2 | 14 | 19 | 1 | 11 | | | |
| | 0.06 ug/mL | 1 | 4 | 6 | 5 | 15 | 19 | 5 | 15 | | | |
| dC7 | 1.5 ug/mL | 17 | 14 | 13 | 1 | 30 | 45 | 4 | 22 | 3.83 | 6.01 | 21.87 |
| | 0.5 ug/mL | 14 | 12 | 4 | 0 | 23 | 28 | 7 | 17 | | | |
| | 0.17ug/mL | 6 | 7 | 0 | 0 | 5 | 10 | 4 | 17 | | | |
| | 0.06 ug/mL | 7 | 6 | 0 | 2 | 5 | 4 | 5 | 11 | | | |
| dC8 | 1.5 ug/mL | 4 | 15 | 12 | 1 | 8 | 14 | 11 | 5 | 0.67 | 0.82 | 3.12 |
| | 0.5 ug/mL | 0 | 4 | 16 | 2 | 8 | 18 | 2 | 5 | | | |
| | 0.17ug/mL | 1 | 2 | 9 | 0 | 4 | 19 | 3 | 1 | | | |
| | 0.06 ug/mL | 1 | 5 | 7 | 0 | 4 | 11 | 8 | 2 | | | |
| dC9 | 1.5 ug/mL | 0 | 17 | 0 | 0 | 17 | 5 | 7 | 44 | 0.33 | 0.52 | 1.88 |
| | 0.5 ug/mL | 4 | 12 | 0 | 0 | 12 | 9 | 1 | 36 | | | |
| | 0.17ug/mL | 4 | 9 | 1 | 0 | 6 | 4 | 1 | 17 | | | |
| | 0.06 ug/mL | 2 | 8 | 0 | 0 | 7 | 1 | 3 | 12 | | | |

**C** SARS-CoV-2 Peptide Pools

| Code | pC1 | pC2 | pC3 | pC4 | pC5 | pC6 | pC7 | pC8 |
|---|---|---|---|---|---|---|---|---|
| **Protein name** | AP3A | NCAP | Nsp12 | Nsp5 | Spike Pool A | Spike Pool B | S-RBD | VME1 |
| **Protein-ID** | P0DTC3 | P0DTC9 | P0DTC9:4393-5324 | P0DTC1:3264-3569 | P0DTC2 | | P0DTC2:319-541 | P0DTC5 |

S. Table 3. Positive scoring of SARS-CoV-2 mega peptide pool-induced SFUs in Pre-COVID Era Subjects (IDs dP1-20) and COVID-Recovered Subjects (dC1-9) using medium control wells (A) or irrelevant peptide pools (B) as the negative control to establish background noise. The raw data for establishing the media control (Medium Controls), and the irrelevant peptide pool control (PP Neg. Contr.) counts are derived from Table 1. SARS-Cov2 peptide pools, decoded in (C), were tested in the specified 4 concentrations on each PBMC sample. In (A) SFU counts are highlighted that exceed the mean plus 3 SD of the four medium control wells as the cut-off for positivity. In (B) positive SFU counts are highlighted taking the mean and 3 SD of the six irrelevant peptide pool-induced SFU counts as the cut-off for positivity. Note: in pre-COVID donors (dP1-18) the SARS-CoV-2-peptide triggered, allegedly coronavirus cross-reactive responses, are diminished once the background is set to filter out noise created by chance cross-reactivity triggered by the irrelevant control peptide pools.

### A SARS-CoV-2 PCR-verified Subjects

| | | EBV Peptide Pools | | | | | | | | | | | | | | PP Neg. Ctrl. | | |
|---|---|---|---|---|---|---|---|---|---|---|---|---|---|---|---|---|---|---|
| ID. | [ ] | EP1 | EP2 | EP3 | EP4 | EP5 | EP6 | EP7 | EP8 | EP9 | EP10 | EP11 | EP12 | EP13 | EP14 | $\bar{x}$ | $\sigma$ | $\bar{x}+3\sigma$ |
| dC1 | 1.5 ug/mL | 2 | 0 | 8 | 19 | 22 | 15 | 9 | 31 | 20 | 18 | 13 | 21 | 1 | 19 | 0.67 | 1.21 | 4.30 |
| | 0.5 ug/mL | 1 | 1 | 5 | 10 | 21 | 11 | 17 | 21 | 12 | 11 | 12 | 35 | 0 | 27 | | | |
| | 0.17ug/mL | 1 | 2 | 0 | 3 | 8 | 14 | 14 | 21 | 13 | 9 | 12 | 32 | 6 | 19 | | | |
| | 0.06 ug/mL | 0 | 1 | 5 | 2 | 13 | 8 | 7 | 20 | 11 | 9 | 9 | 16 | 10 | 23 | | | |
| dC2 | 1.5 ug/mL | 2 | 2 | 1 | 1 | 11 | 13 | 9 | 11 | 6 | 1 | 50 | 13 | 5 | 12 | 1.00 | 1.55 | 5.65 |
| | 0.5 ug/mL | 2 | 2 | 1 | 0 | 6 | 16 | 7 | 8 | 4 | 4 | 26 | 3 | 4 | 14 | | | |
| | 0.17ug/mL | 1 | 2 | 1 | 0 | 3 | 2 | 4 | 10 | 2 | 0 | 10 | 2 | 2 | 10 | | | |
| | 0.06 ug/mL | 0 | 1 | 3 | 1 | 2 | 1 | 1 | 8 | 3 | 1 | 3 | 2 | 0 | 7 | | | |
| dC3 | 1.5 ug/mL | 0 | 0 | 2 | 40 | 18 | 41 | 55 | 2 | 15 | 11 | 57 | 7 | 3 | 13 | 1.17 | 1.17 | 4.67 |
| | 0.5 ug/mL | 1 | 3 | 0 | 24 | 12 | 40 | 45 | 1 | 34 | 4 | 46 | 9 | 3 | 18 | | | |
| | 0.17ug/mL | 0 | 0 | 2 | 23 | 15 | 23 | 39 | 0 | 26 | 4 | 44 | 2 | 4 | 7 | | | |
| | 0.06 ug/mL | 1 | 1 | 1 | 28 | 2 | 20 | 32 | 1 | 17 | 3 | 23 | 0 | 2 | 2 | | | |
| dC4 | 1.5 ug/mL | 1 | 1 | 4 | 51 | 10 | 58 | 41 | 7 | 58 | 5 | 7 | 20 | 0 | 18 | 1.17 | 0.75 | 3.42 |
| | 0.5 ug/mL | 2 | 1 | 1 | 34 | 10 | 96 | 46 | 8 | 65 | 4 | 7 | 19 | 4 | 11 | | | |
| | 0.17ug/mL | 0 | 0 | 0 | 36 | 7 | 73 | 40 | 5 | 70 | 2 | 6 | 11 | 3 | 16 | | | |
| | 0.06 ug/mL | 1 | 4 | 0 | 19 | 2 | 45 | 44 | 7 | 44 | 1 | 6 | 4 | 2 | 13 | | | |
| dC5 | 1.5 ug/mL | 7 | 4 | 9 | 1 | 4 | 10 | 17 | 9 | 33 | 8 | 49 | 5 | 11 | 9 | 3.67 | 3.14 | 13.09 |
| | 0.5 ug/mL | 0 | 4 | 13 | 5 | 5 | 5 | 16 | 19 | 30 | 9 | 20 | 6 | 7 | 3 | | | |
| | 0.17ug/mL | 1 | 0 | 11 | 3 | 4 | 6 | 10 | 9 | 8 | 4 | 18 | 7 | 4 | 8 | | | |
| | 0.06 ug/mL | 0 | 2 | 6 | 2 | 3 | 2 | 5 | 10 | 7 | 7 | 8 | 1 | 5 | 1 | | | |
| dC6 | 1.5 ug/mL | 17 | 4 | 25 | 6 | 5 | 9 | 4 | 3 | 8 | 6 | 10 | 29 | 4 | 4 | 2.50 | 1.22 | 6.17 |
| | 0.5 ug/mL | 6 | 6 | 16 | 3 | 5 | 2 | 8 | 6 | 4 | 5 | 4 | 38 | 6 | 4 | | | |
| | 0.17ug/mL | 4 | 8 | 0 | 2 | 5 | 1 | 16 | 4 | 5 | 3 | 9 | 8 | 9 | 4 | | | |
| | 0.06 ug/mL | 15 | 9 | 2 | 0 | 1 | 2 | 7 | 3 | 4 | 8 | 4 | 3 | 5 | 2 | | | |
| dC7 | 1.5 ug/mL | 10 | 6 | 22 | 4 | 27 | 14 | 9 | 7 | 6 | 6 | 72 | 52 | 2 | 10 | 3.83 | 6.01 | 21.87 |
| | 0.5 ug/mL | 7 | 3 | 6 | 1 | 8 | 3 | 5 | 2 | 1 | 1 | 61 | 19 | 3 | 15 | | | |
| | 0.17ug/mL | 1 | 4 | 0 | 1 | 2 | 7 | 2 | 2 | 1 | 4 | 32 | 4 | 4 | 11 | | | |
| | 0.06 ug/mL | 1 | 1 | 1 | 2 | 3 | 1 | 1 | 6 | 1 | 3 | 17 | 7 | 0 | 5 | | | |
| dC8 | 1.5 ug/mL | 3 | 10 | 4 | 7 | 134 | 9 | 8 | 22 | 8 | 23 | 4 | 0 | 0 | 23 | 0.67 | 0.82 | 3.12 |
| | 0.5 ug/mL | 5 | 6 | 9 | 7 | 103 | 7 | 12 | 22 | 1 | 7 | 1 | 0 | 4 | 12 | | | |
| | 0.17ug/mL | 3 | 6 | 11 | 6 | 130 | 4 | 7 | 12 | 1 | 10 | 3 | 6 | 0 | 5 | | | |
| | 0.06 ug/mL | 6 | 3 | 6 | 8 | 99 | 3 | 2 | 22 | 3 | 4 | 6 | 0 | 1 | 6 | | | |
| dC9 | 1.5 ug/mL | 1 | 23 | 48 | 10 | 7 | 12 | 89 | 25 | 4 | 5 | 75 | 3 | 0 | 9 | 0.33 | 0.52 | 1.88 |
| | 0.5 ug/mL | 1 | 2 | 40 | 2 | 6 | 5 | 49 | 17 | 3 | 2 | 47 | 0 | 0 | 8 | | | |
| | 0.17ug/mL | 0 | 3 | 47 | 7 | 2 | 7 | 36 | 13 | 1 | 7 | 35 | 0 | 2 | 10 | | | |
| | 0.06 ug/mL | 0 | 2 | 50 | 5 | 4 | 8 | 17 | 13 | 0 | 0 | 15 | 0 | 1 | 8 | | | |

### B Pre-COVID Era Subjects

| | | EBV Peptide Pools | | | | | | | | | | | | | | PP Neg. Ctrl. | | |
|---|---|---|---|---|---|---|---|---|---|---|---|---|---|---|---|---|---|---|
| ID. | [ ] | EP1 | EP2 | EP3 | EP4 | EP5 | EP6 | EP7 | EP8 | EP9 | EP10 | EP11 | EP12 | EP13 | EP14 | $\bar{x}$ | $\sigma$ | $\bar{x}+3\sigma$ |
| dP1 | 1.5 ug/mL | 19 | 2 | 2 | 5 | 5 | 30 | 7 | 13 | 30 | 5 | 7 | 9 | 2 | 3 | 2.83 | 1.33 | 6.82 |
| | 0.5 ug/mL | 5 | 2 | 3 | 0 | 0 | 12 | 6 | 8 | 18 | 1 | 4 | 4 | 2 | 5 | | | |
| | 0.17ug/mL | 2 | 2 | 3 | 4 | 1 | 2 | 4 | 6 | 13 | 2 | 3 | 6 | 6 | 2 | | | |
| | 0.06 ug/mL | 0 | 2 | 6 | 4 | 0 | 3 | 5 | 4 | 6 | 2 | 3 | 2 | 2 | 2 | | | |
| dP2 | 1.5 ug/mL | 3 | 1 | 1 | 0 | 2 | 18 | 4 | 2 | 10 | 3 | 2 | 3 | 1 | 6 | 1.83 | 1.17 | 5.34 |
| | 0.5 ug/mL | 0 | 2 | 0 | 1 | 0 | 10 | 4 | 4 | 10 | 2 | 4 | 2 | 1 | 2 | | | |
| | 0.17ug/mL | 1 | 0 | 0 | 2 | 0 | 7 | 2 | 4 | 9 | 2 | 4 | 1 | 0 | 1 | | | |
| | 0.06 ug/mL | 0 | 2 | 1 | 1 | 0 | 8 | 2 | 3 | 4 | 0 | 2 | 1 | 1 | 0 | | | |
| dP3 | 1.5 ug/mL | 5 | 2 | 1 | 88 | 4 | 7 | 1 | 1 | 6 | 3 | 1 | 7 | 3 | 2 | 0.50 | 0.84 | 3.01 |
| | 0.5 ug/mL | 12 | 1 | 2 | 49 | 2 | 10 | 0 | 4 | 4 | 1 | 3 | 6 | 4 | 2 | | | |
| | 0.17ug/mL | 3 | 1 | 1 | 25 | 1 | 4 | 1 | 5 | 2 | 1 | 1 | 0 | 14 | 1 | | | |
| | 0.06 ug/mL | 2 | 1 | 3 | 14 | 1 | 3 | 0 | 1 | 3 | 1 | 1 | 5 | 6 | 0 | | | |
| dP4 | 1.5 ug/mL | 13 | 14 | 7 | 5 | 6 | 11 | 5 | 8 | 35 | 6 | 3 | 20 | 8 | 4 | 2.17 | 2.56 | 9.85 |
| | 0.5 ug/mL | 11 | 7 | 9 | 14 | 14 | 11 | 7 | 7 | 28 | 0 | 13 | 14 | 32 | 19 | | | |
| | 0.17ug/mL | 10 | 4 | 3 | 8 | 7 | 6 | 10 | 4 | 7 | 1 | 8 | 3 | 29 | 13 | | | |
| | 0.06 ug/mL | 7 | 4 | 3 | 8 | 12 | 5 | 1 | 4 | 4 | 2 | 1 | 2 | 3 | 3 | | | |
| dP5 | 1.5 ug/mL | 5 | 3 | 4 | 6 | 2 | 22 | 13 | 18 | 20 | 9 | 14 | 7 | 1 | 6 | 4.33 | 3.56 | 15.01 |
| | 0.5 ug/mL | 2 | 4 | 3 | 5 | 0 | 5 | 11 | 30 | 7 | 6 | 8 | 1 | 2 | 9 | | | |
| | 0.17ug/mL | 2 | 3 | 18 | 5 | 1 | 5 | 16 | 29 | 6 | 5 | 4 | 4 | 1 | 5 | | | |
| | 0.06 ug/mL | 1 | 4 | 3 | 3 | 1 | 1 | 3 | 23 | 5 | 7 | 5 | 2 | 0 | 1 | | | |
| dP6 | 1.5 ug/mL | 6 | 3 | 6 | 3 | 0 | 6 | 4 | 1 | 1 | 6 | 7 | 2 | 2 | 3 | 6.50 | 2.07 | 12.72 |
| | 0.5 ug/mL | 6 | 4 | 7 | 3 | 2 | 7 | 6 | 2 | 6 | 10 | 3 | 5 | 4 | 5 | | | |
| | 0.17ug/mL | 6 | 5 | 11 | 11 | 4 | 2 | 2 | 2 | 5 | 3 | 5 | 5 | 2 | 2 | | | |
| | 0.06 ug/mL | 5 | 6 | 8 | 9 | 0 | 1 | 4 | 3 | 4 | 0 | 3 | 0 | 6 | 1 | | | |
| dP7 | 1.5 ug/mL | 1 | 3 | 3 | 8 | 19 | 58 | 60 | 19 | 77 | 10 | 42 | 12 | 1 | 8 | 1.67 | 1.37 | 5.77 |
| | 0.5 ug/mL | 0 | 2 | 3 | 12 | 14 | 59 | 46 | 18 | 73 | 2 | 27 | 2 | 2 | 5 | | | |
| | 0.17ug/mL | 0 | 1 | 2 | 8 | 9 | 36 | 44 | 10 | 42 | 4 | 10 | 6 | 2 | 2 | | | |
| | 0.06 ug/mL | 1 | 6 | 3 | 5 | 4 | 37 | 36 | 9 | 37 | 1 | 5 | 2 | 0 | 9 | | | |
| dP8 | 1.5 ug/mL | 7 | 1 | 6 | 10 | 10 | 25 | 50 | 14 | 30 | 19 | 17 | 6 | 5 | 24 | 2.50 | 1.05 | 5.65 |
| | 0.5 ug/mL | 11 | 8 | 5 | 11 | 4 | 17 | 36 | 11 | 20 | 12 | 24 | 7 | 7 | 22 | | | |
| | 0.17ug/mL | 5 | 5 | 6 | 11 | 5 | 6 | 22 | 11 | 13 | 14 | 11 | 6 | 5 | 23 | | | |
| | 0.06 ug/mL | 4 | 4 | 13 | 8 | 7 | 9 | 18 | 8 | 9 | 14 | 21 | 2 | 4 | 12 | | | |
| dP9 | 1.5 ug/mL | 1 | 6 | 42 | 5 | 4 | 37 | 21 | 2 | 42 | 0 | 29 | 2 | 0 | 19 | 3.33 | 2.73 | 11.53 |
| | 0.5 ug/mL | 4 | 5 | 45 | 3 | 11 | 81 | 22 | 1 | 26 | 1 | 38 | 0 | 1 | 8 | | | |
| | 0.17ug/mL | 2 | 0 | 37 | 0 | 7 | 68 | 36 | 3 | 26 | 1 | 26 | 3 | 0 | 14 | | | |
| | 0.06 ug/mL | 1 | 1 | 37 | 0 | 5 | 26 | 15 | 2 | 11 | 0 | 10 | 0 | 0 | 5 | | | |
| dP10 | 1.5 ug/mL | 1 | 2 | 1 | 96 | 11 | 230 | 29 | 110 | 222 | 93 | 18 | 12 | 2 | 16 | 4.50 | 10.05 | 34.66 |
| | 0.5 ug/mL | 2 | 3 | 4 | 67 | 1 | 217 | 22 | 91 | 196 | 64 | 12 | 1 | 2 | 6 | | | |
| | 0.17ug/mL | 1 | 2 | 9 | 24 | 5 | 210 | 21 | 76 | 184 | 60 | 7 | 0 | 3 | 4 | | | |
| | 0.06 ug/mL | 0 | 3 | 2 | 12 | 6 | 171 | 19 | 99 | 157 | 40 | 17 | 2 | 3 | 0 | | | |
| dP11 | 1.5 ug/mL | 4 | 52 | 6 | 8 | 143 | 87 | 5 | 10 | 13 | 7 | 9 | 10 | 4 | 10 | 1.50 | 1.87 | 7.11 |
| | 0.5 ug/mL | 5 | 52 | 3 | 4 | 129 | 33 | 10 | 5 | 2 | 4 | 7 | 5 | 1 | 3 | | | |
| | 0.17ug/mL | 1 | 38 | 3 | 1 | 150 | 8 | 6 | 11 | 10 | 5 | 2 | 1 | 1 | 8 | | | |
| | 0.06 ug/mL | 1 | 20 | 3 | 0 | 114 | 3 | 13 | 8 | 3 | 2 | 2 | 0 | 0 | 3 | | | |
| dP12 | 1.5 ug/mL | 0 | 1 | 5 | 7 | 45 | 292 | 83 | 15 | 282 | 6 | 6 | 22 | 2 | 4 | 4.00 | 7.87 | 27.62 |
| | 0.5 ug/mL | 0 | 1 | 4 | 1 | 30 | 274 | 76 | 15 | 249 | 8 | 17 | 16 | 0 | 4 | | | |
| | 0.17ug/mL | 2 | 2 | 4 | 0 | 14 | 250 | 64 | 9 | 254 | 8 | 3 | 7 | 1 | 8 | | | |
| | 0.06 ug/mL | 0 | 2 | 2 | 0 | 1 | 191 | 44 | 2 | 183 | 3 | 6 | 3 | 0 | 4 | | | |
| dP13 | 1.5 ug/mL | 2 | 3 | 1 | 1 | 4 | 33 | 3 | 21 | 19 | 7 | 49 | 10 | 5 | 3 | 2.00 | 1.90 | 7.69 |
| | 0.5 ug/mL | 3 | 1 | 2 | 2 | 4 | 19 | 2 | 33 | 9 | 17 | 38 | 7 | 4 | 2 | | | |
| | 0.17ug/mL | 1 | 2 | 2 | 1 | 3 | 9 | 0 | 23 | 20 | 20 | 29 | 2 | 0 | 0 | | | |
| | 0.06 ug/mL | 3 | 2 | 2 | 2 | 2 | 5 | 1 | 18 | 10 | 7 | 12 | 1 | 3 | 0 | | | |
| dP14 | 1.5 ug/mL | 12 | 2 | 109 | 108 | 6 | 87 | 21 | 70 | 25 | 101 | 153 | 7 | 3 | 9 | 7.33 | 1.75 | 12.59 |
| | 0.5 ug/mL | 8 | 4 | 103 | 82 | 5 | 48 | 16 | 66 | 9 | 103 | 98 | 4 | 3 | 2 | | | |
| | 0.17ug/mL | 2 | 3 | 91 | 47 | 7 | 27 | 48 | 59 | 8 | 110 | 88 | 6 | 2 | 4 | | | |
| | 0.06 ug/mL | 5 | 10 | 88 | 24 | 2 | 13 | 16 | 51 | 7 | 76 | 51 | 3 | 4 | 4 | | | |
| dP15 | 1.5 ug/mL | 5 | 9 | 6 | 16 | 14 | 28 | 16 | 46 | 59 | 23 | 17 | 18 | 6 | 10 | 4.83 | 1.60 | 9.64 |
| | 0.5 ug/mL | 4 | 3 | 5 | 13 | 5 | 8 | 15 | 47 | 20 | 11 | 23 | 13 | 2 | 9 | | | |
| | 0.17ug/mL | 5 | 3 | 5 | 5 | 4 | 2 | 16 | 65 | 7 | 12 | 22 | 7 | 3 | 3 | | | |
| | 0.06 ug/mL | 0 | 3 | 2 | 5 | 3 | 7 | 18 | 40 | 7 | 15 | 9 | 6 | 2 | 5 | | | |
| dP16 | 1.5 ug/mL | 29 | 12 | 11 | 27 | 183 | 94 | 53 | 32 | 15 | 41 | 82 | 22 | 21 | 23 | 12.00 | 4.56 | 25.68 |
| | 0.5 ug/mL | 18 | 15 | 9 | 14 | 87 | 84 | 46 | 34 | 10 | 15 | 90 | 13 | 13 | 11 | | | |
| | 0.17ug/mL | 13 | 18 | 8 | 7 | 40 | 34 | 50 | 24 | 4 | 8 | 49 | 22 | 5 | 14 | | | |
| | 0.06 ug/mL | 8 | 13 | 5 | 10 | 27 | 23 | 48 | 20 | 1 | 11 | 41 | 8 | 10 | 9 | | | |
| dP17 | 1.5 ug/mL | 6 | 4 | 11 | 1 | 70 | 14 | 18 | 1 | 37 | 1 | 3 | 13 | 3 | 4 | 2.50 | 1.22 | 6.17 |
| | 0.5 ug/mL | 2 | 2 | 5 | 2 | 65 | 1 | 11 | 5 | 19 | 21 | 4 | 9 | 4 | 8 | | | |
| | 0.17ug/mL | 1 | 1 | 6 | 0 | 43 | 8 | 8 | 2 | 9 | 1 | 7 | 2 | 0 | 2 | | | |
| | 0.06 ug/mL | 5 | 4 | 6 | 2 | 19 | 6 | 1 | 1 | 2 | 1 | 0 | 3 | 3 | 0 | | | |
| dP18 | 1.5 ug/mL | 4 | 8 | 7 | 3 | 0 | 2 | 9 | 6 | 18 | 1 | 30 | 0 | 4 | 1 | 9.67 | 5.09 | 24.92 |
| | 0.5 ug/mL | 11 | 6 | 16 | 3 | 3 | 0 | 7 | 11 | 12 | 3 | 9 | 0 | 2 | 1 | | | |
| | 0.17ug/mL | 5 | 4 | 8 | 8 | 1 | 3 | 3 | 7 | 1 | 0 | 8 | 0 | 0 | 0 | | | |
| | 0.06 ug/mL | 5 | 2 | 5 | 9 | 0 | 0 | 0 | 3 | 0 | 1 | 2 | 0 | 1 | 0 | | | |

### C EBV Peptide Pools

| Code | Protein name | Protein-ID | # of Peptides |
|---|---|---|---|
| pE1 | BARF1 | P03228 | 53 |
| pE2 | BMLF1 | Q04360 | 117 |
| pE3 | BMRF1 | P03191 | 99 |
| pE4 | BRLF1 | P03209 | 149 |
| pE5 | BZLF1 | P03206 | 59 |
| pE6 | EBNA-LP | Q8AZK7 | 124 |
| pE7 | EBNA1 | P03211 | 158 |
| pE8 | EBNA2 | P12978 | 19 |
| pE9 | EBNA3a | P12977 | 234 |
| pE10 | EBNA3b | Q1HVG4 | 279 |
| pE11 | EBNA3c | Q69140 | 265 |
| pE12 | GP350/340 | P03200 | 224 |
| pE13 | LMP1 | PO3230 | 94 |
| pE14 | LMP2 | P13285 | 122 |

S. Table 4: Affinity analysis of SFU counts triggered by EBV mega peptide pools in PBMC of donors with SARS-CoV-2-PCR-verified infection (A) and in Pre-Covid Era Subjects (B) with the peptides specified in (C). Affinity levels are color-coded. Red: high affinity, defined as four consecutive peptide dilutions eliciting a positive recall response with SFU counts exceeding 3 SD of the negative peptide pool-based background (PP Neg Contr), as specified for each PBMC sample on the right. Orange: intermediate affinity, defined as three consecutive peptide dilutions eliciting a positive recall response. Yellow: low affinity, with only the two highest peptide concentrations eliciting positive SFU counts.

.
